# Measuring genetic differentiation from Pool-seq data

**DOI:** 10.1101/282400

**Authors:** Valentin Hivert, Raphël Leblois, Eric J. Petit, Mathieu Gautier, Renaud Vitalis

## Abstract

The advent of high throughput sequencing and genotyping technologies enables the comparison of patterns of polymorphisms at a very large number of markers. While the characterization of genetic structure from individual sequencing data remains expensive for many non-model species, it has been shown that sequencing pools of individual DNAs (Pool-seq) represents an attractive and cost-effective alternative. However, analyzing sequence read counts from a DNA pool instead of individual genotypes raises statistical challenges in deriving correct estimates of genetic differentiation. In this article, we provide a method-of-moments estimator of *F*_ST_ for Pool-seq data, based on an analysis-of-variance framework. We show, by means of simulations, that this new estimator is unbiased, and outperforms previously proposed estimators. We evaluate the robustness of our estimator to model misspecification, such as sequencing errors and uneven contributions of individual DNAs to the pools. Finally, by reanalyzing published Pool-seq data of different ecotypes of the prickly sculpin *Cottus asper*, we show how the use of an unbiased *F*_ST_ estimator may question the interpretation of population structure inferred from previous analyses.

## INTRODUCTION

It has long been recognized that the subdivision of species into subpopulations, social groups and families fosters genetic differentiation (Wahlund 1928; Wright 1931). Characterizing genetic differentiation as a means to infer unknown population structure is therefore fundamental to population genetics, and finds applications in multiple domains, including conservation biology, invasion biology, association mapping and forensics, among many others. In the late 1940s and early 1950s, Malécot (1948) and Wright (1951) introduced F-statistics to partition genetic variation within and between groups of individuals (Holsinger and Weir 2009; Bhatia et al. 2013). Since then, the estimation of F-statistics has become standard practice (see, e.g., Weir 1996; Weir and Hill 2002; Weir 2012), and the most commonly used estimators of *F*_ST_ have been developed in an analysis-of-variance framework (Cockerham 1969, 1973; Weir and Cockerham 1984), which can be recast in terms of probabilities of identity of pairs of homologous genes (Cockerham and Weir 1987; Rousset 2007; Weir and Goudet 2017).

Assuming that molecular markers are neutral, estimates of *F*_ST_ are typically used to quantify genetic structure in natural populations, which is then interpreted as the result of demographic history (Holsinger and Weir 2009): large *F*_ST_ values are expected for small populations among which dispersal is limited (Wright 1951), or between populations that have long diverged in isolation from each other (Reynolds et al. 1983); when dispersal is spatially restricted, a positive relationship between *F*_ST_ and the geographical distance for pairs of populations generally holds (Slatkin 1993; Rousset 1997). It has also been proposed to characterize the heterogeneity of *F*_ST_ estimates across markers for identifying loci that are targeted by selection (Cavalli-Sforza 1966; Lewontin and Krakauer 1973; Beaumont and Nichols 1996; Vitalis et al. 2001; Akey et al. 2002; Beaumont 2005; Weir et al. 2005; Lotterhos and Whitlock 2014, 2015; Whitlock and Lotterhos 2015).

Next-generation sequencing (NGS) technologies provide unprecedented amounts of polymorphism data in both model and non-model species (Ellegren 2014). Although the sequencing strategy initially involved individually tagged samples in humans (The International HapMap Consortium 2005), whole-genome sequencing of pools of individuals (Pool-seq) is being increasingly popular for population genomic studies (Schlötterer et al. 2014). Because it consists in sequencing libraries of pooled DNA samples and does not require individual tagging of sequences, Pool-seq provides genome-wide polymorphism data at considerably lower cost than sequencing of individuals (Schlötterer et al. 2014). However, non-equimolar amounts of DNA from all individuals in a pool and stochastic variation in the amplification efficiency of individual DNAs have raised concerns with respect to the accuracy of the so-obtained allele frequency estimates, particularly at low sequencing depth and with small pool sizes (Cutler and Jensen 2010; Ellegren 2014; Anderson et al. 2014). Nonetheless, it has been shown that, at equal sequencing effort, Pool-seq provides similar, if not more accurate, allele frequency estimates than individual-based analyses (Futschik and Schlötterer 2010; Gautier et al. 2013). The problem is different for diversity and differentiation parameters, which depend on second moments of allele frequencies or, equivalently, on pairwise measures of genetic identity. With Pool-seq data, however, it is impossible to distinguish pairs of reads that are identical because they were sequenced from a single gene, from pairs of reads that are identical because they were sequenced from two distinct genes that are identical in state (IIS) (Ferretti et al. 2013).

Appropriate estimators of diversity and differentiation parameters must therefore be sought, to account for both the sampling of individual genes from the pool and the sampling of reads from these genes. There has been several attempts to define estimators for the parameter *F*_ST_ for Pool-seq data (Kofler et al. 2011; Ferretti et al. 2013), from ratios of heterozygosities (or from probabilities of genetic identity between pairs of reads) within and between pools. In the following, we will argue that these estimators are biased (i.e., they do not converge towards the expected value of the parameter), and that some of them have undesired statistical properties (i.e., the bias depends upon sample size and coverage). Here, following Cockerham (1969), Cockerham (1973), Weir and Cockerham (1984), Weir (1996), Weir and Hill (2002) and Rousset (2007), we define a method-of-moments estimator of the parameter *F*_ST_ using an analysis-of-variance framework. We then evaluate the accuracy and the precision of this estimator, based on the analysis of simulated datasets, and compare it to estimates defined in the software package PoPoolation2 (Kofler et al. 2011), and in Ferretti et al. (2013). Furthermore, we test the robustness of our estimators to model misspecifications (including unequal contributions of individuals in pools, and sequencing errors). Finally, we reanalyze the prickly sculpin (*Cottus asper*) Pool-seq data (published by Dennenmoser et al. 2017), and show how the use of biased *F*_ST_ estimators in previous analyses may challenge the interpretation of population structure.

Note that throughout this article, we use the term “gene” to designate a segregating genetic unit (in the sense of the “Mendelian gene” from Orgogozo et al. 2016). We further use the term “read” in a narrow sense, as a sequenced copy of a gene. For the sake of simplicity, we will use the term “Ind-seq” to refer to analyses based on individual data in which we further assume that individual genotypes are called without error.

## MODEL

*F*-statistics may be described as intra-class correlations for the probability of identity in state (IIS) of pairs of genes (Cockerham and Weir 1987; Rousset 1996, 2007), and *F*_ST_ is best defined as:

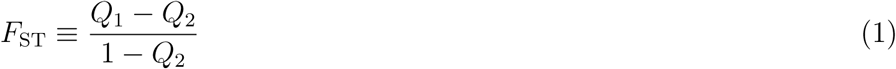

where *Q*_1_ is the IIS probability for genes sampled within subpopulations, and *Q*_2_ is the IIS probability for genes sampled between subpopulations. In the following, we develop an estimator of *F*_ST_ for Pool-seq data, by decomposing the total variance of gene frequencies in an analysis-of-variance framework. A complete derivation of the model is provided in the Supplemental File S1.

For the sake of clarity, the notation used throughout this article is given in Table 1. We first derive our model for a single locus, and eventually provide a multilocus estimator of *F*_ST_. Consider a sample of *n*_d_ subpopulations, each of which is made of *n*_i_ genes (*i* = 1,…,*n_d_*) sequenced in pools (hence *n*_i_ is the haploid sample size of the *i*th pool). We define *c_ij_* as the number of reads sequenced from gene *j* (*j = 1,…,n*_i_) in subpopulation *i* at the locus considered. Note that *c_ij_* is a latent variable, that cannot be directly observed from the data. Let *X_ijr:k_* be an indicator variable for read *r* (*r* = 1,…, *c_ij_*) from gene *j* in subpopulation *i*, such that *X_ijr:k_* = 1 if the rth read from the *j*th gene in the *i*th deme is of type *k*, and *X_ijr:k_* = 0 otherwise. In the following, we use standard dot notations for sample averages, i.e.: 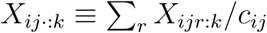, 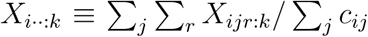 and 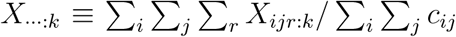. The analysis of variance is based on the computation of sums of squares, as follows:

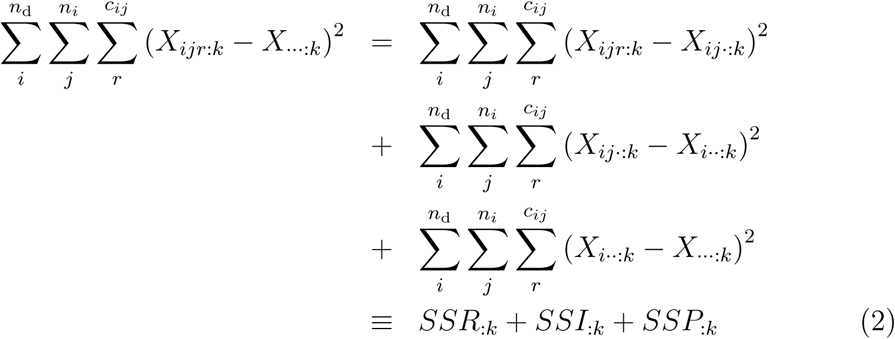

**Table 1.** Summary of main notations.

| Notation | Parameter definition |
| --- | --- |
| $X_{ijr:k}$ | Indicator variable: $X_{ijr:k} = 1$ if the $r$th read from the $j$th individual in the $i$th pool is of type $k$, and $X_{ijr:k} = 0$ otherwise |
| $r_{i:k} = \sum_j \sum_r X_{ijr:k}$ | Number of reads of type $k$ in the $i$th pool |
| $c_{ij}$ | Number of reads sequenced from individual $j$ in sub-population $i$ (unobserved individual coverage) |
| $C_{1i} \equiv \sum_j c_{ij}$ | Total number of reads in the $i$th pool (pool coverage) |
| $C_1 \equiv \sum_i C_{1i}$ | Total number of reads in the full sample (total coverage) |
| $C_2 \equiv \sum_i C_{1i}^2$ | Squared number of reads in the full sample |
| $n_i$ | Total number of genes the $i$th pool (haploid pool size) |
| $y_{i:k}$ | (Unobserved) number of genes of type $k$ in the $i$th pool |
| $\pi_k \equiv \mathbb{E}(X_{ijr:k})$ | Expected frequency of reads of type $k$ in the full sample |
| $\hat{\pi}_{ij:k} \equiv X_{ij\cdot:k}$ | (Unobserved) average frequency of reads of type $k$ for individual $j$ in the $i$th pool |
| $\hat{\pi}_{i:k} \equiv X_{i\cdot\cdot:k}$ | Average frequency of reads of type $k$ in the $i$th pool |
| $\hat{\pi}_k \equiv X_{\dots:k}$ | Average frequency of reads of type $k$ in the full sample |
| $Q_1$ (resp. $Q_2$) | IIS probability for two genes sampled within (resp. between) pools |
| $Q_1^r$ (resp. $Q_2^r$) | IIS probability for two reads sampled within (resp. between) pools |
| $\hat{Q}_1^{\text{pool}}$ (resp. $\hat{Q}_2^{\text{pool}}$) | Unbiased estimator of the IIS probability for genes sampled within (resp. between) populations |

As is shown in the Supplemental File S1, the expected sums of squares depend on the expectation of the allele frequency *π_k_* over all replicate populations sharing the same evolutionary history, as well as on the IIS probability *Q*_1:_*_k_* that two genes in the same pool are both of type *k*, and the IIS probability *Q*_2:_*_k_* that two genes from different pools are both of type k. Taking expectations (see the detailed computations in the Supplemental File S1), one has:

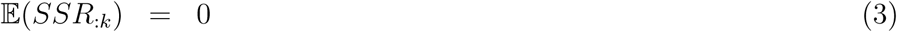

for reads within individual genes, since we assume that there is no sequencing error, i.e. all the reads sequenced from a single gene are identical and *X_ijr:k_* = *X_ij·:k_* for all r. For reads between genes within pools, we get:

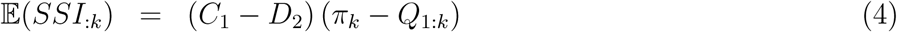

where 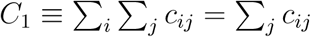 is the total number of reads in the full sample (total coverage), *C*_1_*_i_* is the coverage of the *i*th pool and 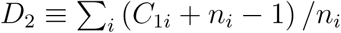. *D*_2_ arises from the assumption that the distribution of the read counts *c_ij_*· is multinomial (i.e., that all genes contribute equally to the pool of reads; see Equation A15 in Supplemental File S1). For reads between genes from different pools, we have:

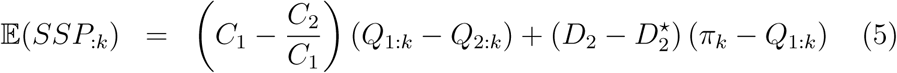

where 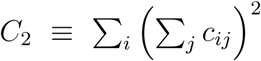 and 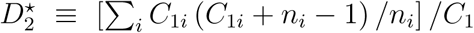 (see Equation A16 in Supplemental File S1). Rearranging Equations 4-5, and summing over alleles, we get:

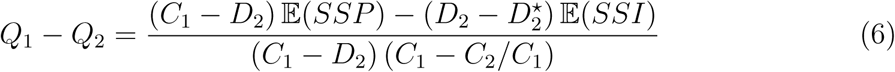

and:

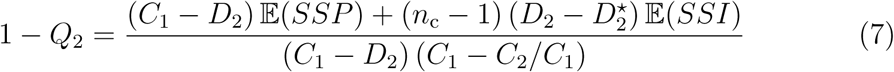

where 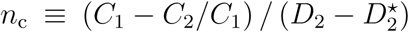. Let *MSI* = *SSI*/ (*C*_1_ − *D*_2_) and 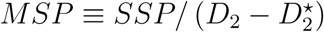. Then:

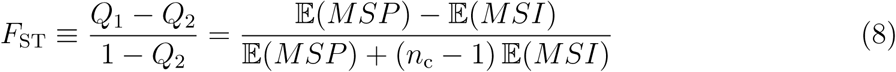

which yields the method-of-moments estimator:

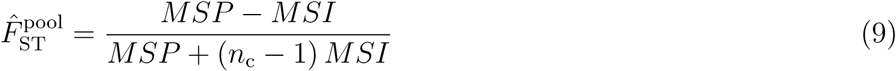

where

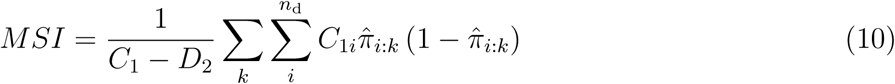

and:

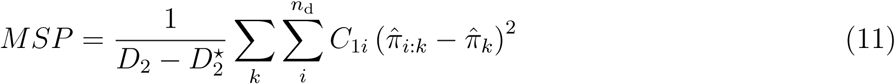

(see Equations A25 and A26 in Supplemental File S1). In Equations 10 and 11, 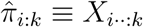 is the average frequency of reads of type *k* within the *i*th pool, and 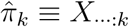 is the average frequency of reads of type *k* in the full sample. Note that from the definition of 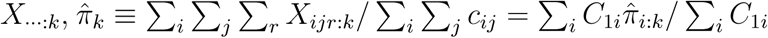 is the weighted average of the sample frequencies with weights equal to the pool coverage. This is equivalent to the weighted analysis-of-variance in Cockerham (1973) (see also Weir and Cockerham 1984; Weir 1996; Weir and Hill 2002; Rousset 2007; Weir and Goudet 2017). Finally, the full expression of 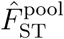 in terms of sample frequencies reads:

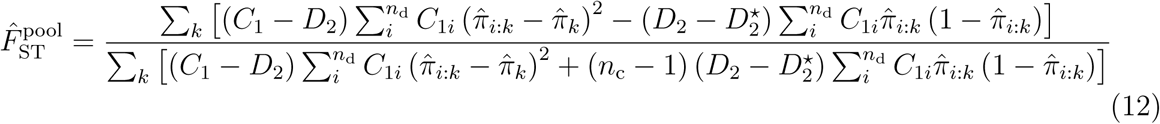

If we take the limit case where each gene is sequenced exactly once, we recover the Ind-seq model: assuming *c_ij_* = 1 for all (*i,j*), then 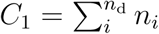, 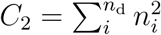, *D*_2_ = *n*_d_ and 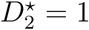. Therefore, *n*_c_ = (*C*_1_ − *C*_2_/*C*_1_) / (*n*_d_ *−* 1), and Equation 9 reduces exactly to the estimator of *F*_ST_ for haploids: see Weir (1996), p. 182, and Rousset (2007), p. 977.

As in Reynolds et al. (1983), Weir and Cockerham (1984), Weir (1996) and Rousset (2007), a multilocus estimate is derived as the sum of locus specific numerators over the sum of locus-specific denominators:

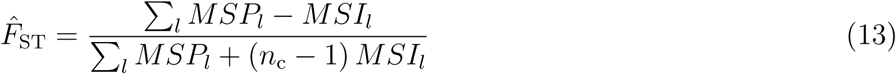

where *MSI* and *MSP* are subscripted with *l* to denote the *l*th locus. For Ind-seq data, Bhatia et al. (2013) refer to this multilocus estimate as a “ratio of averages” by opposition to an “average of ratios”, which would consist in averaging single-locus *F*_ST_ over loci. This approach is justified in the Appendix of Weir and Cockerham (1984) and in Bhatia et al. (2013), who analyzed both estimates by means of coalescent simulations. Note that Equation 13 assumes that the pool size is equal across loci. Also note that the construction of the estimator in Equation 13 is different from Weir and Cockerham’s (1984). These authors defined their multilocus estimator as a ratio of sums of components of variance (a, *b* and c in their notation) over loci, which give the same weight to all loci, whatever the number of sampled genes at each locus. Equation 13 follows Genepop’s rationale (Rousset 2008), which gives instead more weight to loci that are more intensively covered.

## MATERIALS AND METHODS

### Simulation study

*Generating individual genotypes:* we first generated individual genotypes using `ms` (Hudson 2002), assuming an island model of population structure (Wright 1931). For each simulated scenario, we considered 8 demes, each made of *N* = 5, 000 haploid individuals. The migration rate (*m*) was fixed to achieve the desired value of *F*_ST_ (0.05 or 0.2), using Equation 6 in Rousset (1996) leading, e.g., to *M* ≡ *2Nm* = 16.569 for *F*_ST_ = 0.05 and *M* = 3.489 for *F*_ST_ = 0.20. The mutation rate was set at *μ* = 10^-6^, giving *θ* ≡ 2*N μ* = 0.01. We considered either fixed, or variable sample sizes across demes. In the later case, the haploid sample size *n* was drawn independently for each deme from a Gaussian distribution with mean 100 and standard deviation 30; this number was rounded up to the nearest integer, with min. 20 and max. 300 haploids per deme. We generated a very large number of sequences for each scenario, and sampled independent single nucleotide polymorphisms (SNPs) from sequences with a single segregating site. Each scenario was replicated 50 times (500 times for Figures 3 and S2).

*Pool sequencing:* for each `ms` simulated dataset, we generated Pool-seq data by drawing reads from a binomial distribution (Gautier et al. 2013). More precisely, we assume that for each SNP, the number *r_i:k_* of reads of allelic type *k* in pool *i* follows:

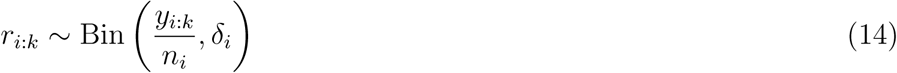

where *y_i:k_* is the number of genes of type *k* in the *i*th pool, *n_i_* is the total number of genes in pool *i* (haploid pool size), and *δ_i_* is the simulated total coverage for pool *i*. In the following, we either consider a fixed coverage, with *δ_i_* = ∆ for all pools and loci, or a varying coverage across pools and loci, with *δ_i_* ~ Pois(∆).

*Sequencing error:* we simulated sequencing errors occurring at rate *μ*_e_ = 0.001, which is typical of Illumina sequencers (Glenn 2011; Ross et al. 2013). We assumed that each sequencing error modifies the allelic type of a read to one of three other possible states with equal probability (there are therefore four allelic types in total, corresponding to four nucleotides). Note that only biallelic markers are retained in the final datasets. Also note that, since we initiated this procedure with polymorphic markers only, we neglect sequencing errors that would create spurious SNPs from monomorphic sites. However, such SNPs should be rare in real datasets, since markers with a low minimum read count (MRC) are generally filtered out.

*Experimental error:* non-equimolar amounts of DNA from all individuals in a pool and stochastic variation in the amplification efficiency of individual DNAs are sources of experimental errors in pool sequencing. To simulate experimental errors, we used the model derived by Gautier et al. (2013). In this model, it is assumed that the contribution *η_ij_* = *C_ij_*/*C*_1_*_i_* of each gene *j* to the total coverage of the *i*th pool (*C*_1_*_i_*) follows a Dirichlet distribution:

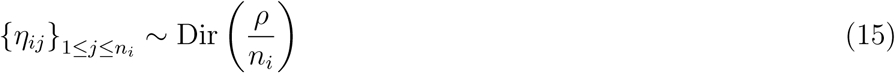

where the parameter *ρ* controls the dispersion of gene contributions around the value *η_ij_* = 1/*n_i_*, expected if all genes contributed equally to the pool of reads. For convenience, we define the experimental error *∊* as the coefficient of variation of %·, i.e.: 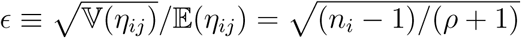 (see Gautier et al. 2013). When *∊* tends toward 0 (or equivalently when *ρ* tends to infinity), all individuals contribute equally to the pool, and there is no experimental error. We tested the robustness of our estimates to values of *∊* comprised between 0.05 and 0.5. The case *∊* = 0.5 could correspond, for example, to a situation where (for *n_i_* = 10) 5 individuals contribute 2.8 × more reads than the other 5 individuals.

### Other estimators

For the sake of clarity, a summary of the notation of the *F*_ST_ estimators used throughout this article is given in Table 2.

**Table 2.** Definition of the *F*_ST_ estimators used in the text.

| Notation | Definition |
| --- | --- |
| $\hat{F}_{ST}^{pool}$ | Equation 9 |
| FRP<sub>13</sub> | Ferretti et al. (2013) and Equations 16,20–21 |
| NC<sub>83</sub> | Nei and Chesser (1983) |
| PP2<sub>d</sub> | Kofler et al. (2011) and Equations 16–18 |
| PP2<sub>a</sub> | Kofler et al. (2011) and Equation 19 |
| WC<sub>84</sub> | Weir and Cockerham (1984) |

PP2_d_ : this estimator of *F*_ST_ is implemented by default in the software package PoPoolation2 (Kofler et al. 2011). It is based on a definition of the parameter *F*_ST_ as the overall reduction in average heterozygosity relative to the total combined population (see, e.g., Nei and Chesser 1983):

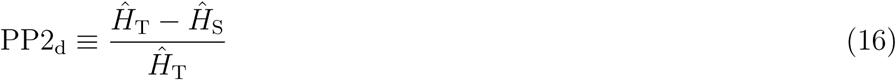

where 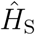 is the average heterozygosity within subpopulations, and 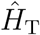is the average heterozygosity in the total population (obtained by pooling together all subpopulation to form a single virtual unit). In PoPoolation2, 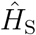 is the unweighted average of within-subpopulation heterozygosities:

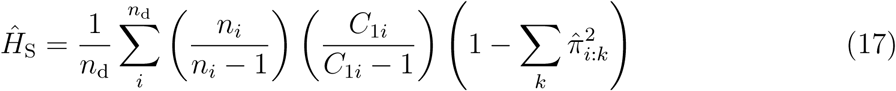

(using the notation from Table 1). Note that in PoPoolation2, PP2_d_ is restricted to the case of two subpopulations only (*n*_d_ = 2). The two ratios in the right-hand side of Equation 17 are presumably borrowed from Nei (1978) to provide an unbiased estimate, although we found no formal justification for the expression in Equation 17 for Pool-seq data. The total heterozygosity is computed as (using the notation from Table 1):

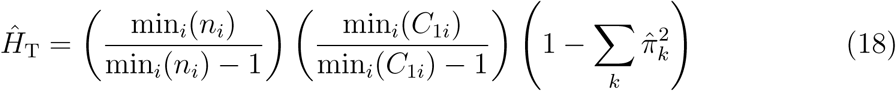

PP2_a_ : this is the alternative estimator of *F*_ST_ provided in the software package PoPoolation2. It is based on an interpretation by Kofler et al. (2011) of Karlsson et al.’s (2007) estimator of *F*_ST_, as:

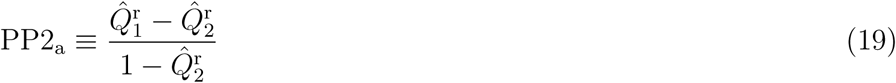

where 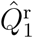 and 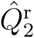 are the frequencies of identical pairs of reads within and between pools, respectively, computed by simple counting of IIS pairs. These are estimates of 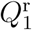, the IIS probability for two reads in the same pool (whether they are sequenced from the same gene or not) and 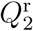, the IIS probability for two reads in different pools. Note that the IIS probabiliy 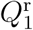 is different from *Q*_1_ in Equation 1, which, from our definition, represents the IIS probability between distinct genes in the same pool. This approach therefore confounds pairs of reads within pools that are identical because they were sequenced from a single gene, from pairs of reads that are identical because they were sequenced from distinct, yet IIS genes.

FRP_13_ : this estimator of *F*_ST_ was developed by Ferretti et al. (2013) (see their Equations 3 and 10-13). Ferretti et al. (2013) use the same definition of *F*_ST_ as in Equation 16 above, although they estimate heterozygosities within and between pools as “average pairwise nucleotide diversities”, which, from their definitions, are formally equivalent to IIS probabilities. In particular, they estimate the average heterozygosity within pools as (using the notation from Table 1):

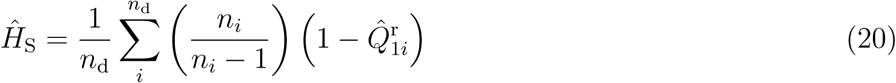

and the total heterozygosity among the *n*_d_ populations as:

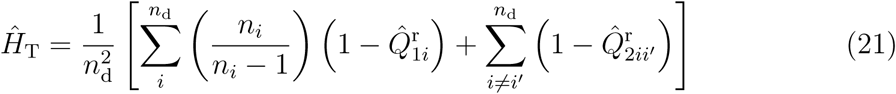

### Analyses of Ind-seq data

For the comparison of Ind-seq and Pool-seq datasets, we computed *F*_ST_ on subsamples of 5,000 loci. These subsamples were defined so that only those loci that were polymorphic in all coverage conditions were retained, and the same loci were used for the analysis of the corresponding Ind-seq data. For the latter, we used either the Nei and Chesser’s (1983) estimator based on a ratio of heterozygosity (see Equation 16 above), hereafter denoted by NC_83_, or the analysis-of-variance estimator developed by Weir and Cockerham (1984), hereafter denoted by WC_84_.

All the estimators were computed using custom functions in the R software environment for statistical computing, version 3.3.1 (R Core Team 2017). All these functions were carefully checked against available software packages, to ensure that they provided strictly identical results.

### Application example: *Cottus asper*

Dennenmoser et al. (2017) investigated the genomic basis of adaption to osmotic conditions in the prickly sculpin (*Cottus asper*), an abundant eury-haline fish in northwestern North America. To do so, they sequenced the whole-genome of pools of individuals from two estuarine populations (CR, Capilano River Estuary; FE, Fraser River Estuary) and two freshwater populations (PI, Pitt Lake and HZ, Hatzic Lake) in southern British Columbia (Canada). We downloaded the four corresponding BAM files from the Dryad Digital Repository (doi: 10.5061/dryad.2qg01) and combined them into a single mpileup file using `SAMtools` version 0.1.19 (Li et al. 2009) with default options, except the maximum depth per BAM that was set to 5,000 reads. The resulting file was further processed using a custom `awk` script, to call SNPs and compute read counts, after discarding bases with a Base Alignment Quality (BAQ) score lower than 25. A position was then considered as a SNP if: (*i*) only two different nucleotides with a read count > 1 were observed (nucleotides with ≤ 1 read being considered as a sequencing error); (*ii*) the coverage was comprised between 10 and 300 in each of the four alignment files; (*iii*) the minor allele frequency, as computed from read counts, was ≥ 0.01 in the four populations. The final data set consisted of 608,879 SNPs.

Our aim here was to compare the population structure inferred from pairwise estimates of *F*_ST_, using the estimator 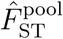 on the one hand, and PP2_d_ on the other hand. Then, to conclude on which of the two estimators performs better, we compared the population structure inferred from 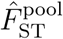 and PP2_d_ to that inferred from the Bayesian hierarchical model implemented in the software package BayPass (Gautier 2015). BayPass allows indeed the robust estimation of the scaled covariance matrix of allele frequencies across populations for Pool-seq data, which is known to be informative about population history (Pickrell and Pritchard 2012). The elements of the estimated matrix can be interpreted as pairwise and population-specific estimates of differentiation (Coop et al. 2010), and therefore provide a comprehensive description of population structure that makes full use of the available data.

### Data availability

The authors state that all data necessary for confirming the conclusions presented in this article are fully represented within the article, figures, and tables. Supplemental Tables S1–S3 and Figures S1–S4 are available at FigShare, along with a complete derivation of the model in the Supplemental File S1 at FigShare.

## RESULTS

### Comparing Ind-seq and Pool-seq estimates of *F*_ST_

Single-locus estimates 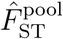 are highly correlated with the classical estimates WC_84_ (Weir and Cockerham 1984) computed on the individual data that were used to generate the pools in our simulations (see Figure 1). The variance of 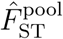 across independent replicates decreases as the coverage increases. The correlation between 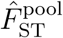 and WC_84_ is stronger for multilocus estimates (see Figure S1A).

**Figure 1.**
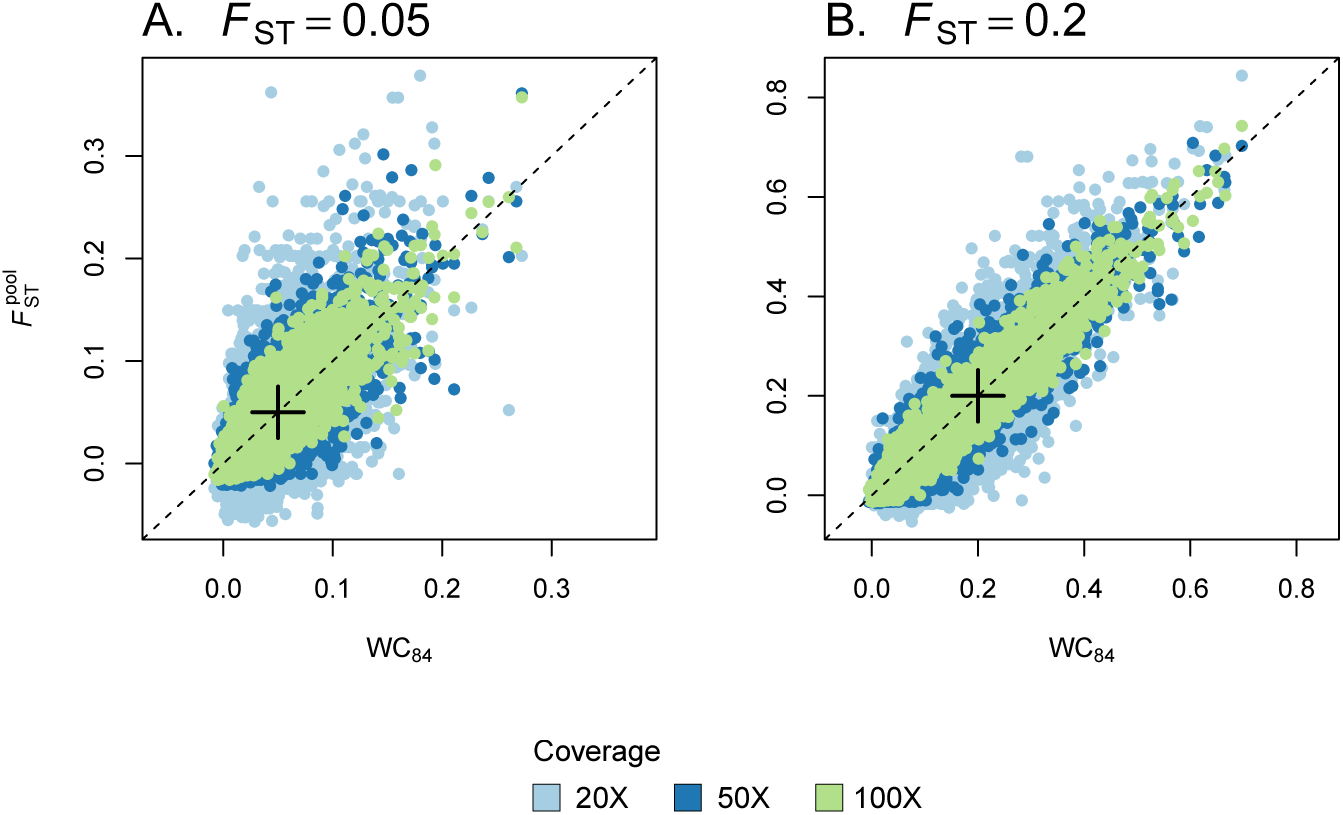
Single-locus estimates of *F*_ST_. We compared single-locus estimates of *F*_ST_ based on allele count data inferred from individual genotypes (Ind-seq), using the WC_84_ estimator, to 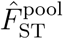 estimates from Pool-seq data. We simulated 5,000 SNPs using `ms` in an island model with *n*_d_ = 8 demes. We used two migration rates corresponding to *F*_ST_ = 0.05 (A) and *F*_ST_ = 0.20 (B). The size of each pool was fixed to 100. We show the results for different coverages (20X, 50X and 100X). In each graph, the cross indicates the simulated value of *F*_ST_.

### Comparing Pool-seq estimators of *F*_ST_

We found that our estimator 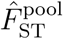 has extremely low bias (< 0.5% over all scenarios tested: see Tables 3 and S1-S3). In other words, the average estimates across multiple loci and replicates closely equals the expected value of the *F*_ST_ parameter, as given by Equation 6 in Rousset (1996), which is based on the computation of IIS probabilities in an island model of population structure. In all the situations examined, the bias did neither depend on the sample size (i.e., the size of each pool) nor on the coverage (see Figure 2). Only the variance of the estimator across independent replicates decreases as the sample size increases and/or as the coverage increases. At high coverage, the mean and root mean squared error (RMSE) of 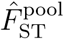 over independent replicates are virtually indistinguishable from that of the WC_84_ estimator (see Table S1).

**Table 3.** Overall *F*_ST_ estimates from multiple pools. Overall *F*_ST_ was estimated for various conditions of expected *F*_ST_, pool size (*n*) and coverage (Cov.). For Pool-seq data, we computed our estimator 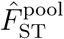 (Equation 13). The mean (RMSE) over 50 independent replicates of the `ms` simulations are provided, for all populations (*n*_d_ = 8). For comparison, we computed WC_84_ from allele count data inferred from individual genotypes (Ind-seq).

| $F_{ST}$ | $n$ | Pool-seq | | Ind-seq |
| --- | --- | --- | --- | --- |
| | | Cov. | $\hat{F}_{ST}^{pool}$ | WC<sub>84</sub> |
| 0.05 | 10 | 20× | 0.050 (0.002) |  |
| 0.05 | 10 | 50× | 0.051 (0.002) | 0.050 (0.002) |
| 0.05 | 10 | 100× | 0.050 (0.002) |  |
| 0.05 | 100 | 20× | 0.050 (0.001) |  |
| 0.05 | 100 | 50× | 0.050 (0.001) | 0.051 (0.001) |
| 0.05 | 100 | 100× | 0.050 (0.001) |  |
| 0.20 | 10 | 20× | 0.200 (0.002) |  |
| 0.20 | 10 | 50× | 0.201 (0.002) | 0.201 (0.002) |
| 0.20 | 10 | 100× | 0.201 (0.002) |  |
| 0.20 | 100 | 20× | 0.201 (0.003) |  |
| 0.20 | 100 | 50× | 0.202 (0.003) | 0.203 (0.003) |
| 0.20 | 100 | 100× | 0.203 (0.003) |  |

**Figure 2.**
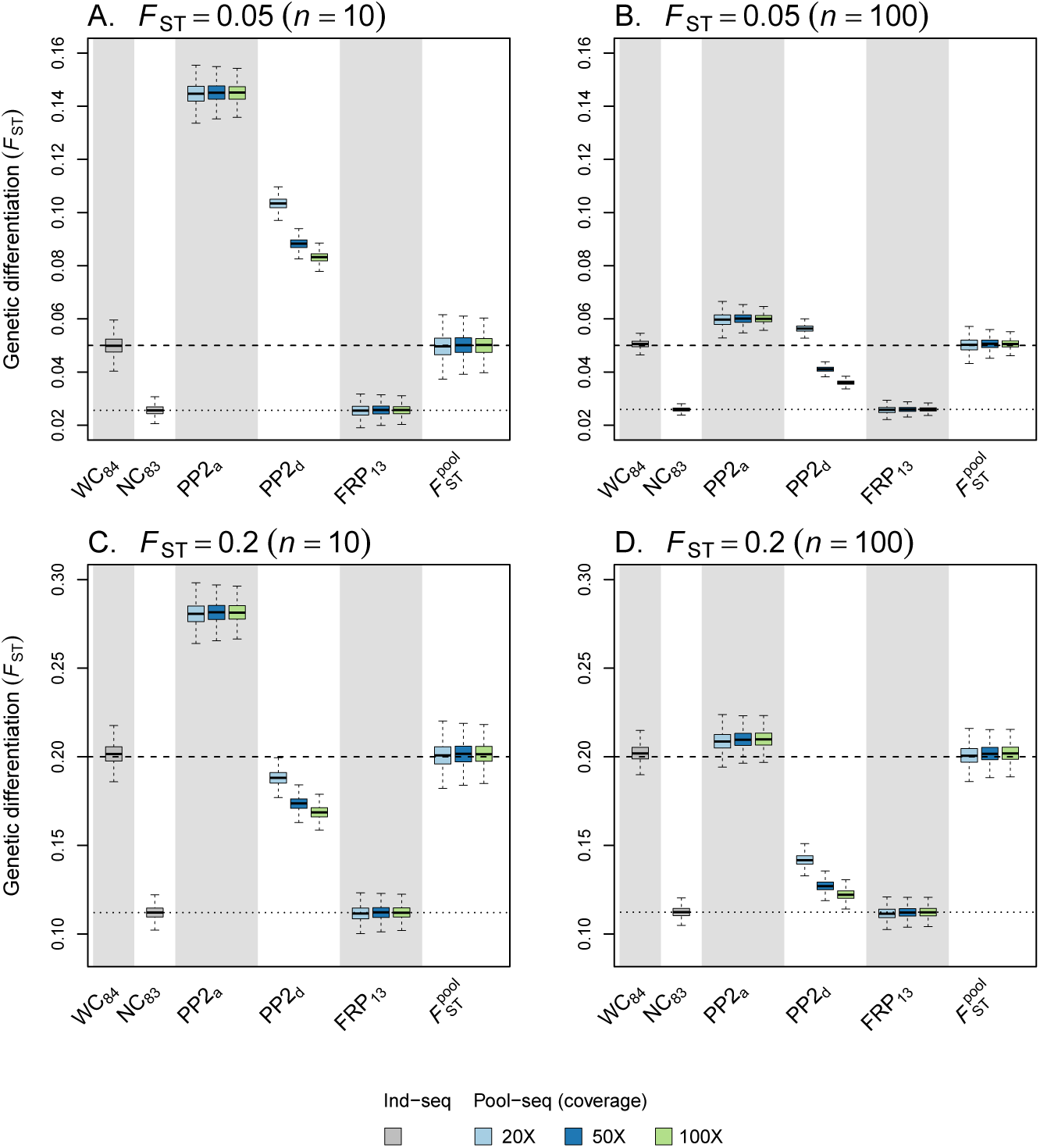
Precision and accuracy of pairwise estimators of *F*_ST_. We considered two estimators based on allele count data inferred from individual genotypes (Ind-seq): WC_84_ and NC_83_. For pooled data, we computed the two estimators implemented in the software package PoPoolation2, that we refer to as PP2_d_ and PP2_a_, as well as the FRP_13_ estimator and our estimator 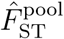 (Equation 13). Each boxplot represents the distribution of multilocus *F*_ST_ estimates across all pairwise comparisons in an island model with *n*_d_ = 8 demes, and across 50 independent replicates of the `ms` simulations. We used two migration rates, corresponding to *F*_ST_ = 0.05 (A-B) or *F*_ST_ = 0.20 (C-D). The size of each pool was either fixed to 10 (A and C) or to 100 (B and D). For Pool-seq data, we show the results for different coverages (20X, 50X and 100X). In each graph, the dashed line indicates the simulated value of *F*_ST_ and the dotted line indicates the median of the distribution of NC_83_ estimates.

Figure 3 shows the RMSE of *F*_ST_ estimates for a wide range of pool sizes and coverage. The RMSE decreases as the pool size and/or the coverage increases. The *F*_ST_ estimates are more precise and accurate when differentiation is low. Figure 3 provides some clues to evaluate the pool size and the coverage that is necessary to achieve the same RMSE than for Ind-seq data. Consider, for example, the case of samples of *n* = 20 haploids. For *F*_ST_ ≤ 0.05 (in the conditions of our simulations), the RMSE of *F*_ST_ estimates based on Pool-seq data tends to the RMSE of *F*_ST_ estimates based on Ind-seq data either by sequencing pools of ca. 200 haploids at 20X, or by sequencing pools of 20 haploids at ca. 200X. However, the same precision and accuracy are achieved by sequencing ca. 50 haploids at ca. 50X.

**Figure 3.**
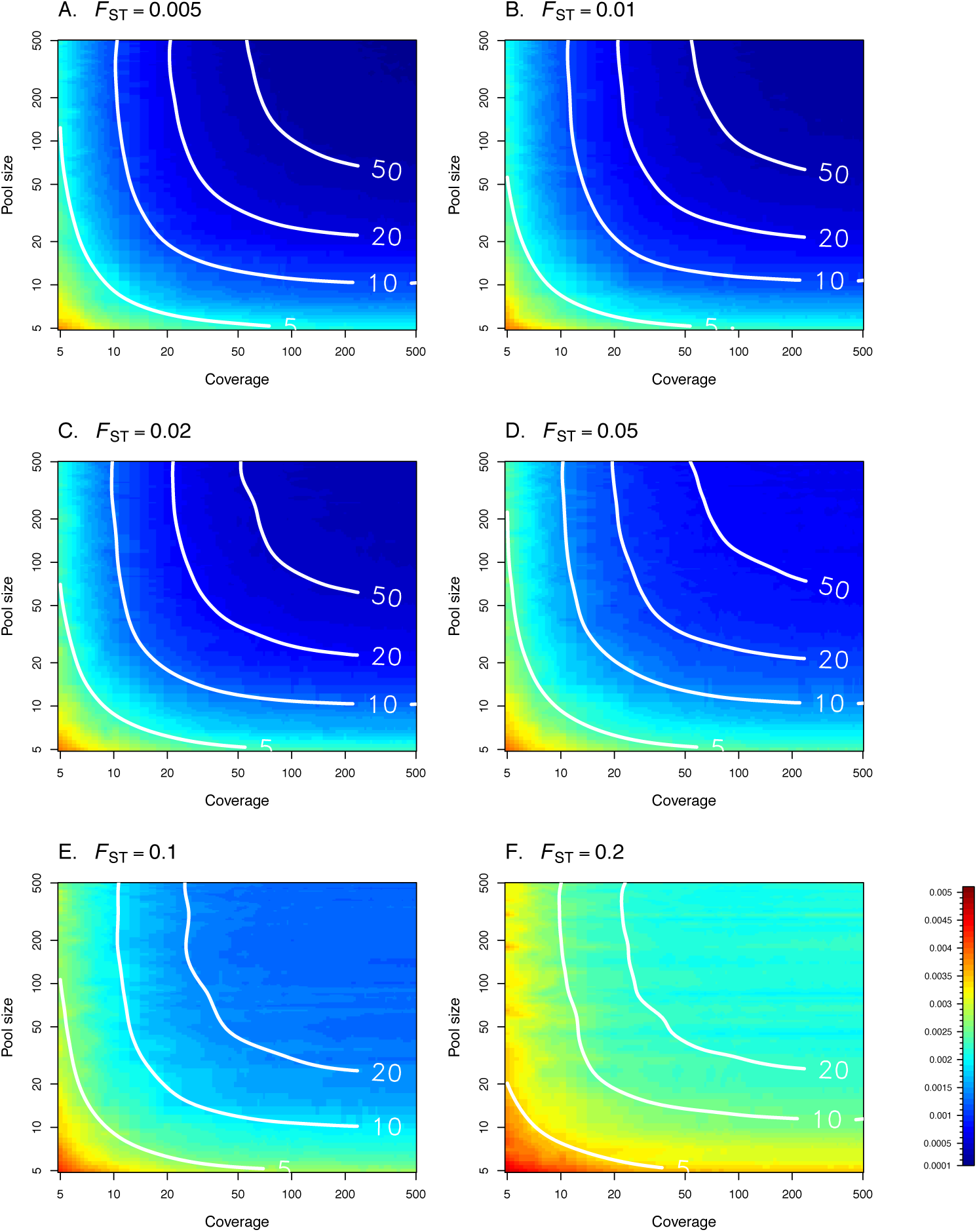
Root mean squared error (RMSE) of *F*_ST_ estimates for a wide range of pool sizes and coverage, with *F*_ST_ varying from 0.005 to 0.2 (A – F). Each density plot gives the RMSE of our estimator 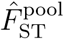, using simple linear interpolation from a set of 44 × 44 pairs of pool size and coverage values. For each pool size and coverage, 500 replicates of 5,000 markers were simulated. Plain white isolines represent the RMSE of the WC_84_ estimator computed from Ind-seq data, for various sample sizes (*n* = 5, 10, 20, and 50). Each isoline was fitted using a thin plate spline regression with smoothing parameter λ = 0.005, implemented in the `fields` package for R (Nychka et al. 2017).

Conversely, we found that PP2_d_ (the default estimator of *F*_ST_ implemented in the software package PoPoolation2) is biased when compared to the expected value of the parameter. We observed that the bias depends on both the sample size, and the coverage (see Figure 2). We note that, as the coverage and the sample size increase, PP2_d_ converges to the estimator NC_83_ (Nei and Chesser 1983) computed from individual data (see Figure S1B). This argument was used by Kofler et al. (2011) to validate the approach, even though the estimates PP2d depart from the true value of the parameter (Figure S1B–C).

The second of the two estimators of *F*_ST_ implemented in PoPoolation2, that we refer to as PP2_a_, is also biased (see Figure 2). We note that the bias decreases as the sample size increases. However, the bias does not depend on the coverage (only the variance over independent replicates does). The estimator developed by Ferretti et al. (2013), that we refer to as FRP_i3_, is also biased (see Figure 2). However, the bias does neither depend on the pool size, nor on the coverage (only the variance over independent replicates does). FRP_i3_ converges to the estimator NC_83_, computed from individual data (see Figure 2). At high coverage, the mean and RMSE over independent replicates are virtually indistinguishable from that of the NC_83_ estimator.

Last, we stress out that our estimator 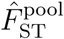 provides estimates for multiple populations, and is therefore not restricted to pairwise analyses, contrary to PoPoolation2’s estimators. We show that, even at low sample size and low coverage, Pool-seq estimates of differentiation are virtually indistinguishable from classical estimates for Ind-seq data (see Table 3).

### Robustness to unbalanced pool sizes and variable sequencing coverage

We evaluated the accuracy and the precision of the estimator 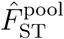 when sample sizes differ across pools, and when the coverage varies across pools and loci (see Figure 4). We found that, at low coverage, unequal sampling or variable coverage causes a negligible departure from the median of WC_84_ estimates computed on individual data, which vanishes as the coverage increases. At 100X coverage, the distribution of 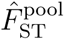 estimates is almost indistinguishable from that of WC_84_ (see Figure 4 and Tables S2–S3).

**Figure 4.**
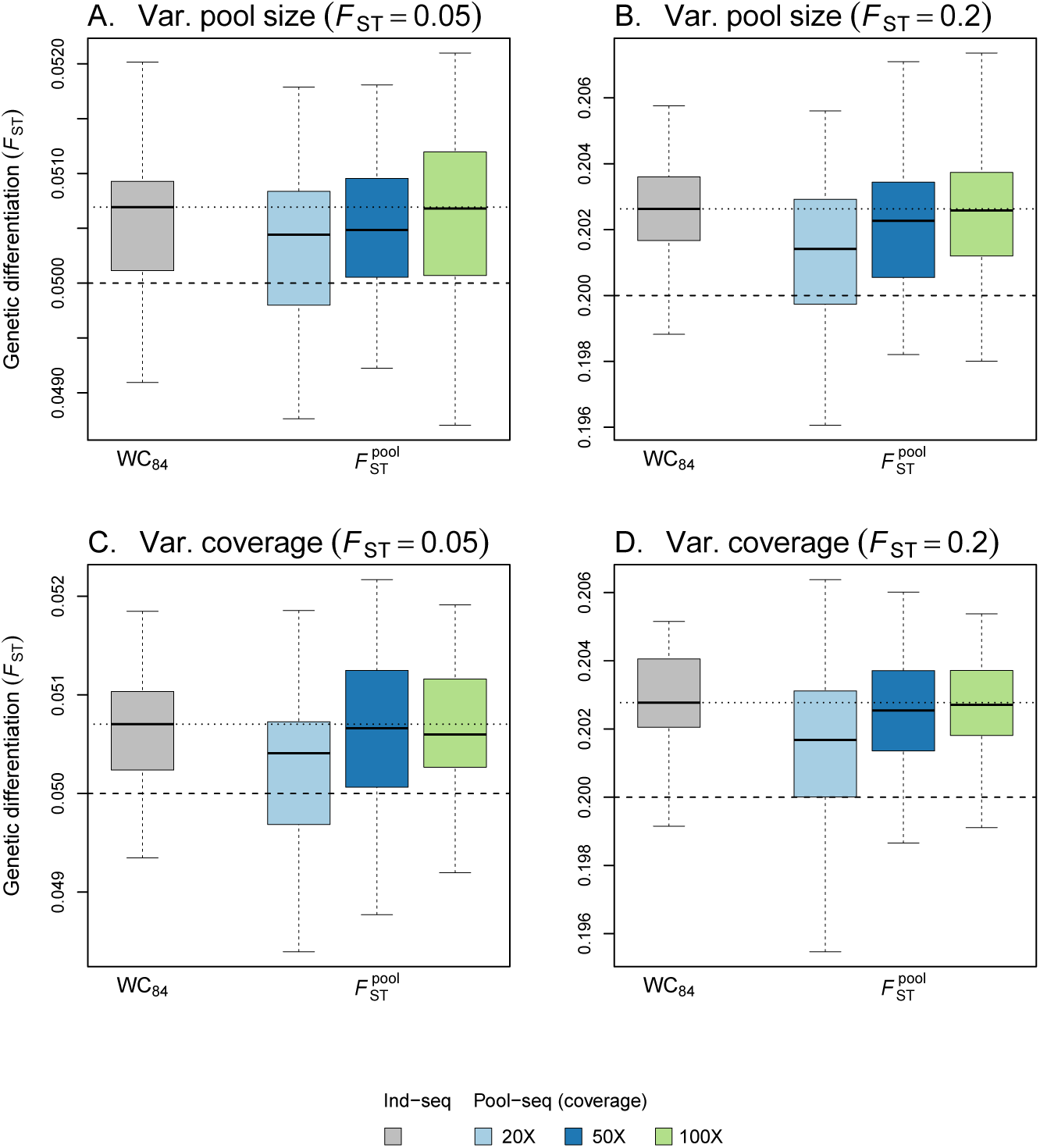
Precision and accuracy of *F*_ST_ estimates with varying pool size or varying coverage. Our estimator 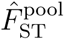 (Equation 13) was calculated from Pool-seq data over all loci and demes and compared to the estimator WC_84_, computed from allele count data inferred from individual genotypes (Ind-seq). Each boxplot represents the distribution of multilocus *F*_ST_ estimates across 50 independent replicates of the `ms` simulations. We used two migration rates, corresponding to *F*_ST_ = 0.05 (A and C) or *F*_ST_ = 0.20 (B and D). In A-B the pool size was variable across demes, with haploid sample size *n* drawn independently for each deme from a Gaussian distribution with mean 100 and standard deviation 30; *n* was rounded up to the nearest integer, with min. 20 and max. 300 haploids per deme. In C-D, the pool size was fixed (*n* = 100), and the coverage (*δ_i_*) was varying across demes and loci, with *δ_i_* ~ Pois(∆) where ∆ ∈ {20, 50,100}. For Pool-seq data, we show the results for different coverages (20X, 50X and 100X). In each graph, the dashed line indicates the simulated value of *F*_ST_ and the dotted line indicates the median of the distribution of WC84 estimates.

### Robustness to sequencing and experimental errors

Figure 5 shows that sequencing errors cause a negligible negative bias for 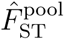 estimates. Filtering (using a minimum read count of 4) improves estimation slightly, but only at high coverage (Figure 6B). It must be noted, though, that filtering increases the bias in the absence of sequencing error, especially at low coverage (Figure 6A). With experimental error, i.e., when individuals do not contribute evenly to the final set of reads, we observed a positive bias for 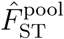 estimates (Figure 5). We note that the bias decreases as the size of the pools increases. Figure S2 shows the RMSE of *F*_ST_ estimates for a wider range of pool sizes, coverage and experimental error rate. For *∊* ≥ 0.25, increasing the coverage cannot improve the quality of the inference, if the pool size is too small. When Pool-seq experiments are prone to large experimental error rates, increasing the size of pools is the only way to improve the estimation of *F*_ST_. Filtering (using a minimum read count of 4) does not improve estimation (Figure 6C).

**Figure 5.**
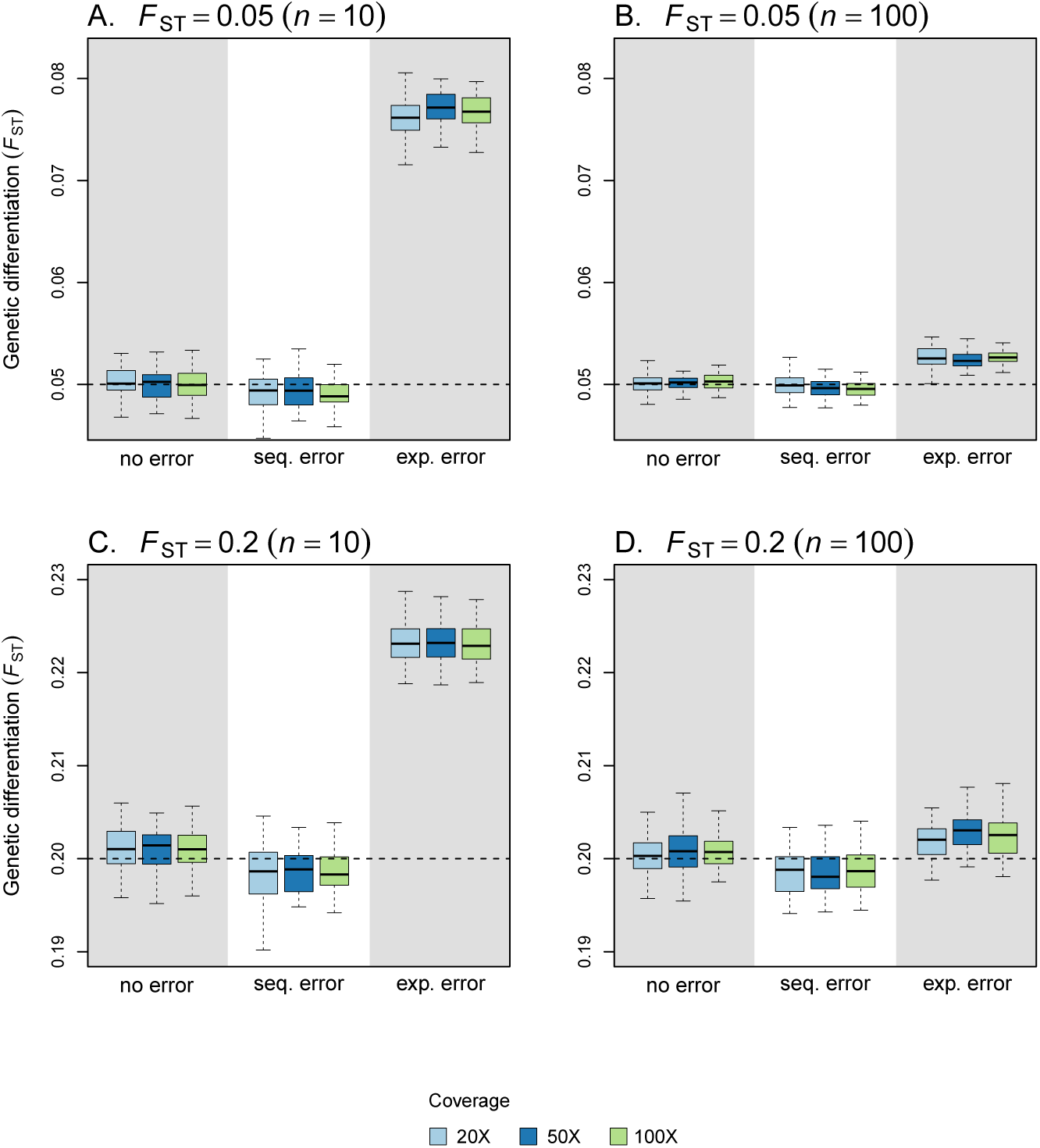
Precision and accuracy of *F*_ST_ estimates with sequencing and experimental errors. Our estimator 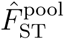 (Equation 13) was computed from Pool-seq data over all loci and demes without error, with sequencing error (occurring at rate *μ*_e_ = 0.001), and with experimental error (∊ = 0.5). Each boxplot represents the distribution of multilocus *F*_ST_ estimates across 50 independent replicates of the `ms` simulations. We used two migration rates, corresponding to *F*_ST_ = 0.05 (A-B) or *F*_ST_ = 0.20 (C-D). The size of each pool was either fixed to 10 (A and C) or to 100 (B and D). For Pool-seq data, we show the results for different coverages (20X, 50X and 100X). In each graph, the dashed line indicates the simulated value of *F*_ST_.

**Figure 6.**
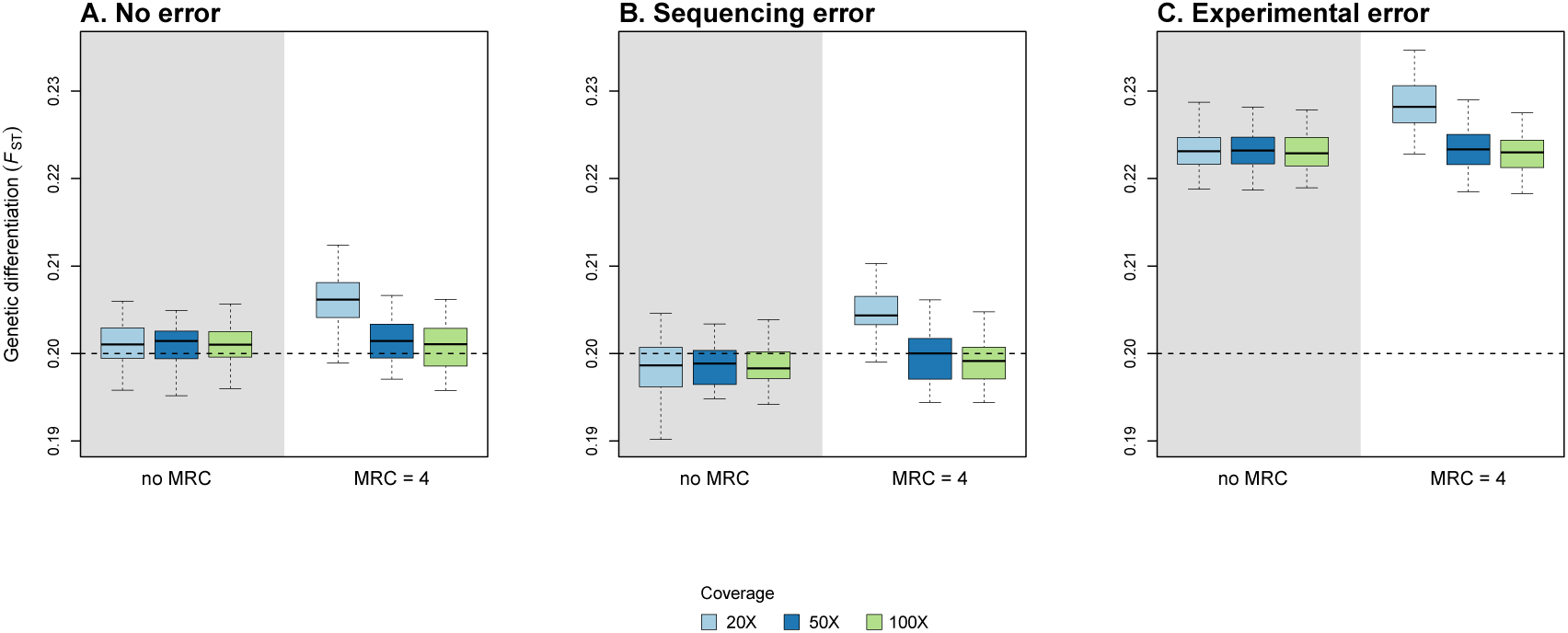
Precision and accuracy of *F*_ST_ estimates with and without filtering. Our estimator 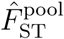 (Equation 13) was computed from Pool-seq data over all loci and demes without error (A), with sequencing error (B) and with experimental error (C) (see the legend of Figure 5 for further details). For each case, we computed *F*_ST_ without filtering (no MRC) and with filtering (using a minimum read count MRC = 4). Each boxplot represents the distribution of multilocus *F*_ST_ estimates across 50 independent replicates of the `ms` simulations. We used a migration rate corresponding to *F*_ST_ = 0.20, and pool size *n* = 10. We show the results for different coverages (20X, 50X and 100X). In each graph, the dashed line indicates the simulated value of *F*_ST_.

### Application example

The reanalysis of the prickly sculpin data revealed larger pairwise estimates of multilocus *F*_ST_ using PP2_d_ estimator, as compared to 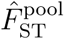 (see Figure 7A). Furthermore, we found that 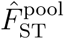 estimates are smaller for within-ecotype pairwise comparisons as compared to between-ecotype comparisons. Therefore, the inferred relationships between samples based on pairwise 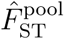 estimates show a clear-cut structure, separating the two estuarine samples from the freshwater ones (see Figure 7C). We did not recover the same structure using PP2_d_ estimates (see Figure 7B). Supportingly, the scaled covariance matrix of allele frequencies across samples is consistent with the structure inferred from 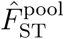 estimates (see Figure 7D).

**Figure 7.**
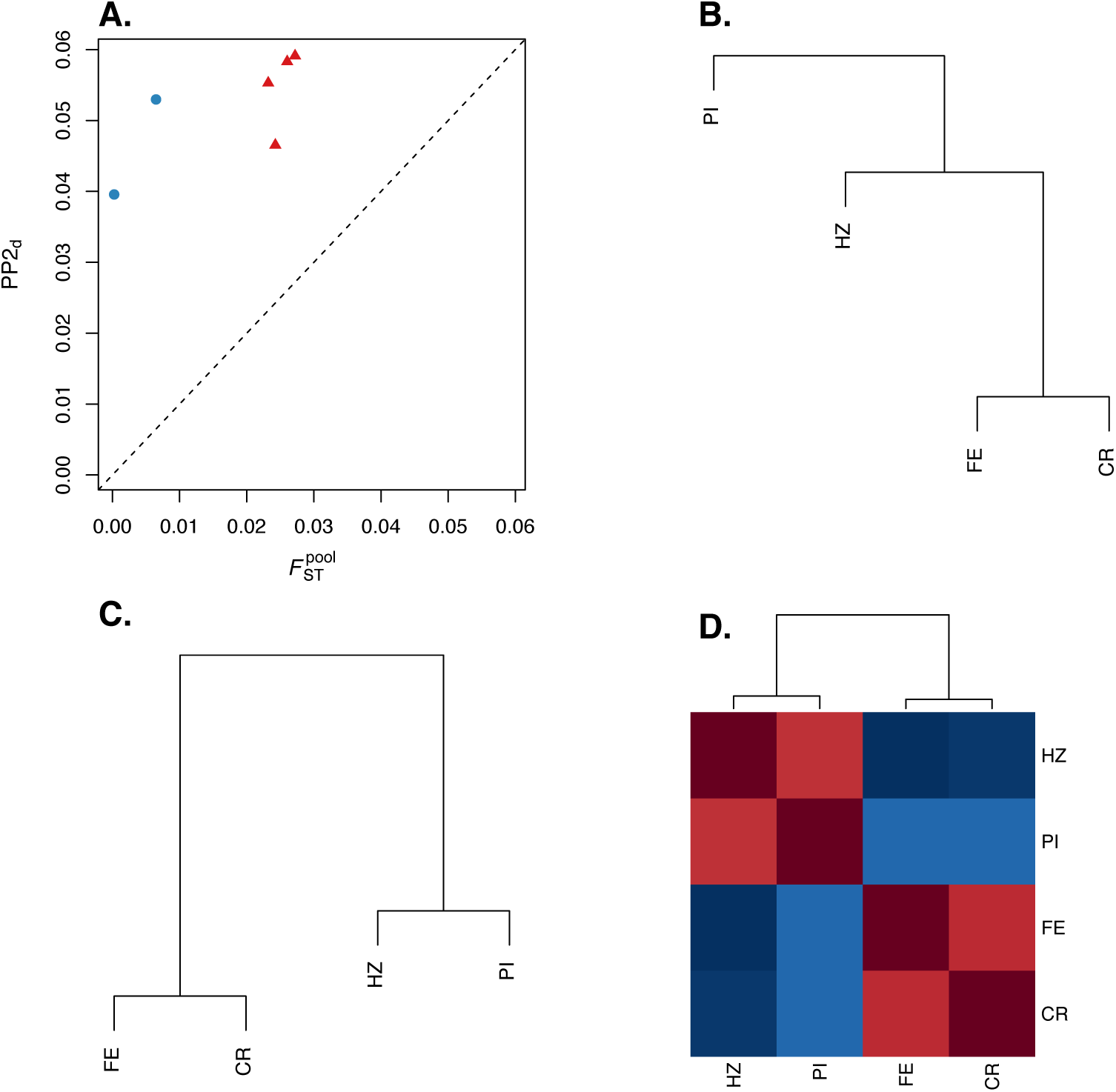
Analysis of the prickly sculpin (*Cottus asper*) Pool-seq data. In (A) we compare the pairwise *F*_ST_ estimates PP2_d_, and 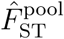 (Equation 13) for all pairs of populations from the estuarine (CR and FE) and freshwater samples (PI and HZ). Within-ecotype comparisons are depicted as blue dots, and between-ecotype comparisons as red triangles. In (B-C) we show a UPGMA hierarchical cluster analyses based on PP2_d_ (B) and 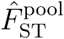 (C) pairwise estimates. In (D), we show a heatmap representation of the scaled covariance matrix among the four *C. asper* populations, inferred from the Bayesian hierarchical model implemented in the software package BayPass.

## DISCUSSION

Whole-genome sequencing of pools of individuals is being increasingly popular for population genomic research on both model and non-model species (Schlötterer et al. 2014). The development of dedicated software packages (reviewed in Schlötterer et al. 2014) has undoubtedly something to do with the breadth of research questions that have been tackled using pool-sequencing. Yet, the analysis of population structure from Pool-seq data is complicated by the double sampling process of genes from the pool and sequence reads from those genes (Ferretti et al. 2013).

The naive approach that consists in computing *F*_ST_ from read counts, as if they were allele counts (e.g., as in Chen et al. 2016), ignores the extra variance brought by the random sampling of reads from the gene pool during Pool-seq experiments. Furthermore, such computation fails to consider the actual number of lineages in the pool (haploid pool size). Altogether, these limits may result in severely biased estimates of differentiation when the pool size is low (see Figure S3). A possible alternative is to compute *F*_ST_ from allele counts imputed from read counts using a maximum-likelihood approach conditional on the haploid size of the pools (e.g., as in Smadja et al. 2012; Leblois et al. 2018), or from allele frequencies estimated using a model-based method that accounts for the sampling effects and the sequencing error probabilities inherent to pooled NGS experiments (see Fariello et al. 2017). However, these latter approaches may only be accurate in situations where the coverage is much larger than pool size, allowing to reduce sampling variance of reads (see Figure S3).

Here, we therefore developed a new estimator of the parameter *F*_ST_ for Pool-seq data, in an analysis-of-variance framework (Cockerham 1969, 1973). The accuracy of this estimator is barely distinguishable from that of the Weir and Cockerham’s (1984) estimator for individual data. Furthermore, does neither depend on the pool size nor on the coverage, and is robust to unequal pool sizes and varying coverage across demes and loci. In our analysis the frequency of reads within pools is a weighted average of the sample frequencies with weights equal to the pool coverage. Therefore, our approach follows Cockerham’s (1973) one, which he referred to as a weighted analysis-of-variance (see also Weir and Cockerham 1984; Weir 1996; Weir and Hill 2002; Weir and Goudet 2017).

With unequal pool sizes, weighted and unweighted analyses differ. As discussed recently in Weir and Goudet (2017), the unweighted approach seems appropriate when the between component exceed the within component, i.e. when *F*_ST_ is large (Tukey 1957). It turns out that optimal weighting depends upon the parameter to be estimated (Cockerham 1973) and is only efficient at lower levels of differentiation (Robertson 1962). In a likelihood analysis of the island model, Rousset (2007) derived asymptotically efficient weights that are proportional to 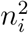for the sum of squares of different samples (i.e., as in Robertson 1962). To the best of our knowledge, such optimal weighting has never been considered in the literature. Nevertheless, if these arguments are true for estimators of variance components, they do not necessarily apply to estimates of intra-class correlations (Cockerham 1973).

### Analysis of variance and probabilities of identity

In the analysis-of-variance framework, *F*_ST_ is defined in Equation 1 as an intraclass correlation for the probability of identity in state (Cockerham and Weir 1987; Rousset 1996). Extensive statistical literature is available on estimators of intraclass correlations. Beside analysis-of-variance estimators, introduced in population genetics by Cockerham (1969, 1973), estimators based on the computation of probabilities of identical response within and between groups have been proposed (see, e.g., Fleiss 1971; Fleiss and Cuzick 1979; Mak 1988; Ridout et al. 1999; Wu et al. 2012), which were originally referred to as kappa-type statistics (Fleiss 1971; Landis and Koch 1977). These estimators have later been endorsed in population genetics, where the “probability of identical response” was then interpreted as the frequency with which the genes are alike (Cockerham 1973; Cockerham and Weir 1987; Weir 1996; Rousset 2007; Weir and Goudet 2017).

This suggests that, with Pool-seq data, another strategy could consist in computing *F*_ST_ from IIS probabilities between (unobserved) pairs of genes, which requires that unbiased estimates of such quantities are derived from read count data. We have done so in the second section of the Supplemental File S1, and we provide alternative estimators of *F*_ST_ for Pool-seq data (see Equations A44 and A48 in Supplemental File S1). These estimators (denoted by 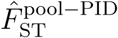 and 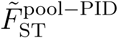) have exactly the same form as the analysis-of-variance estimator if the pools have all the same size and if the number of reads per pool is constant (Equation A33). This echoes the derivations by Rousset (2007) for Ind-seq data, who showed that the analysis-of-variance approach (Weir and Cockerham 1984) and the simple strategy of estimating IIS probabilities by counting identical pairs of genes provide identical estimates when sample sizes are equal (see Equation A28 and also Cockerham and Weir 1987; Weir 1996; Karlsson et al. 2007). With unbalanced samples, we found that analysis-of-variance estimates have better precision and accuracy than IIS-based estimates, particularly for low levels of differentiation (see Figure S4). Interestingly, we found that IIS-based estimates of *F*_ST_ for Pool-seq data have generally lower bias and variance if the overall estimates of IIS probabilities within and between pools are computed as unweighted averages of population-specific or pairwise estimates (see Equations A39 and A43), as compared to weighted averages. Equation A28 further shows that our estimator may be rewritten as a function close to 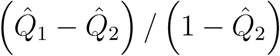, except that it also depends on the sums 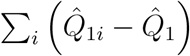 in both the numerator and the denominator. This suggests that if the *Q*_1_*_i_*’s differ among subpopulations, then our estimator provides an estimate of an average of population-specific *F*_ST_ (Weir and Hill 2002; Weir and Goudet 2017).

It follows from the derivations in the Supplemental File S1 that the estimator PP2_a_ (Equation 19) is biased, because the IIS probability between pairs of reads within a pool 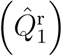 is a biased estimator of the IIS probability between pairs of distinct genes in that pool (see Equation A34 in Supplemental File S1). This is so, because the former confounds pairs of reads that are identical because they were sequenced from a single gene copy, from pairs of reads that are identical because they were sequenced from distinct, yet IIS genes.

A more justified estimator of *F*_ST_ has been proposed by Ferretti et al. (2013), based on previous developments by Futschik and Schlötterer (2010). Note that, although they defined *F*_ST_ as a ratio of functions of heterozygosities, they actually worked with IIS probabilities (see Equations 20 and 21). However, although their Equation 20 is strictly identical to our Equation A34 in Supplemental File S1, we note that they computed the total heterozygosity by integrating over pairs of genes sampled both within and between populations (see Equation 21), which may explain the observed bias (see Figure 2).

### Comparison with alternative estimators

An alternative framework to Weir and Cockerham’s (1984) analysis-of-variance has been developed by Masatoshi Nei and coworkers to estimate *F*_ST_ from gene diversities (Nei 1973, 1977; Nei and Chesser 1983; Nei 1986). The estimator PP2_d_ (see Equations 16-18) implemented in the software package PoPoolation2 (Kofler et al. 2011) follows this logic. However, it has long been recognized that both frameworks are fundamentally different in that the analysis-of-variance approach considers both statistical and genetic (or evolutionary) sampling, whereas Nei and coworkers’ approach do not (Weir and Cockerham 1984; Excoffier 2007; Holsinger and Weir 2009). Furthermore, the expectation of Nei and coworkers’ estimators depend upon the number of sampled populations, with a larger bias for lower numbers of sampled populations (Goudet 1993; Excoffier 2007; Weir and Goudet 2017). This is so, because the computation of the total diversity in Equations 18 and 21 includes the comparison of pairs of genes from the same subpopulation, whereas the computation of IIS probabilities between subpopulations do not (see, e.g., Excoffier 2007). Therefore, we do not recommend using the estimator PP2_d_ implemented in the software package PoPoolation2 (Kofler et al. 2011).

### Applications in evolutionary ecology studies

Pool-seq is being increasingly used in many application domains (Schlötterer et al. 2014), such as conservation genetics (see, e.g., Fuentes-Pardo 2017), invasion biology (see, e.g., Dexter et al. 2018) and evolutionary biology in a broader sense (see, e.g., Collet et al. 2016). These studies use a large range of methods, which aim at characterizing fine-scaled population structure (see, e.g., Fischer et al. 2017), reconstructing past demography (see, e.g., Chen et al. 2016; Leblois et al. 2018), or identifying footprints of natural or artificial selection (see, e.g., Chen et al. 2016; Fariello et al. 2017; Leblois et al. 2018).

Here, we reanalyzed the Pool-seq data produced by Dennenmoser et al. (2017), who investigated the adaptive genomic divergence between freshwater and brackish-water ecotypes of the prickly sculpin *C. asper*, an abundant euryhaline fish in northwestern North America. Measuring pairwise genetic differentiation between samples using 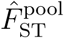, we found a clear-cut structure separating the freshwater from the brackish-water ecotypes. Such genetic strucure supports the hypothesis that populations are locally adaptated to osmotic conditions in these two contrasted habitats, as discussed in Dennenmoser et al. (2017). This structure, which is at odds with that inferred from PP2_d_ estimates, is not only supported by the scaled covariance matrix of allele frequencies, but also by previous microsatellite-based studies, who showed that populations were genetically more differentiated between ecotypes than within ecotypes (Dennenmoser et al. 2014, 2015).

### Limits of the model and perspectives

We have shown that the stronger source of bias for the 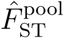 estimate is unequal contributions of individuals in pools. This is so, because we assume in our model that the read counts are multinomially distributed, which supposes that all genes contribute equally to the pool of reads (Gautier et al. 2013), i.e. that there is no variation in DNA yield across individuals and that all genes have equal sequencing coverage (Rode et al. 2018). Because the effect of unequal contribution is expected to be stronger with small pool sizes, it has been recommended to use pool-seq with at least 50 diploid individuals per pool (Lynch et al. 2014; Schlötterer et al. 2014). However, this limit may be overly conservative for allele frequency estimates (Rode et al. 2018), and we have shown here that we can achieve very good precision and accuracy of *F*_ST_ estimates with smaller pool sizes. Furthermore, because genotypic information is lost during Pool-seq experiments, we assume in our derivations that pools are haploid (and therefore that *F*_IS_ is nil). Analyzing non-random mating populations (e.g., in selfing species) is therefore problematic.

Finally, our model, as in Weir and Cockerham (1984), formally assumes that all populations provide independent replicates of some evolutionary process (Excoffier 2007; Holsinger and Weir 2009). This may be unrealistic in many natural populations, which motivated Weir and Hill (2002) to derive a population-specific estimator of *F*_ST_ for Ind-seq data (see also Vitalis et al. 2001). Even though the use of Weir and Hill’s (2002) estimator is still scarce in the literature (but see Weir et al. 2005; Vitalis 2012), Weir and Goudet (2017) recently proposed a re-interpretation of population-specific estimates of *F*_ST_ in terms of allelic matching proportions, which are strictly equivalent to IIS probabilities between pairs of genes. It would therefore be straightforward to extend Weir and Goudet’s (2017) estimator of population-specific *F*_ST_ for the analysis of Pool-seq data, using the unbiased estimates of IIS probabilies provided in the Supplemental File S1.

## DATA ACCESSIBILITY

A R package, called `poolfstat`, which impletements *F*_ST_ estimates for Pool-seq data, is available at the Comprehensive *r* Archive Network (CRAN): https://cran.r-project.org/web/packages/poolfstat/index.html.

## ACKNOWLEDGEMENTS

We thank Alexandre Dehne-Garcia for his assistance in using computer farms. Analyses were performed on the genotoul bioinformatics platform Toulouse Midi-Pyrénées (bioinfo.genotoul.fr) and the CBGP HPC computational platform. This work is part of Valentin Hivert’s Ph.D., who was supported by a grant from the INRA’s Plant Health and Environment (SPE) Division, and by the BiodivERsA project EXOTIC (ANR-13-EBID-0001). Part of this work was supported by the ANR project SWING (ANR-16-CE02-0015) of the French National Research Agency, and by the CORBAM project of the French region Hauts-de-France. We thank two anonymous reviewers for their positive comments and suggestions.

## SUPPLEMENTAL FILE S1: DETAILED MATHEMATICAL DERIVATIONS

### Analysis of variance for Pool-seq data

In the following, we first derive our model for a single locus. Consider a sample of *n*_d_ subpopulations, each of which is made of *n*_i_ genes (*i* = 1,…, *n*_d_) sequenced in pools (hence *n*_i_ is the haploid sample size of the *i*th pool). We define *C_ij_* as the number of reads sequenced from gene *j* (*j* = 1,…, *n*_i_) in subpopulation *i* at the locus considered. Note that *C_ij_* is a latent variable, that cannot be directly observed from the data. Let *X_ijr:k_* be an indicator variable for read *r* (*r* = 1,…, *C_ij_*) from gene *j* in subpopulation *i*, such that *X_ijr:k_* = 1 if the *r*th read from the *j*th gene in the *i*th deme is of type *k*, and *X_ijr:k_* = 0 otherwise. In the following, we use standard dot notations for sample averages, i.e.: 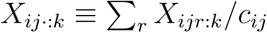, 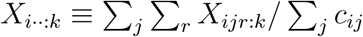 and 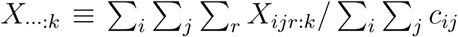. The analysis of variance is based on the computation of sums of squares, as follows:

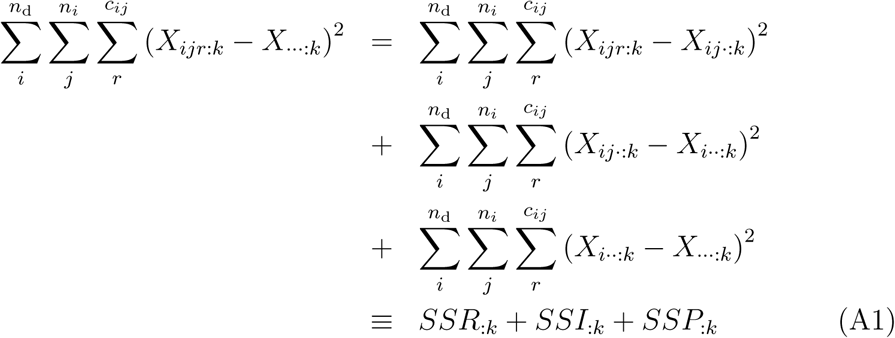

We express the sum of squares for reads within individuals as:

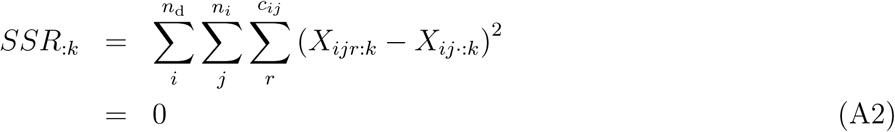

since we assume that there is no sequencing error, i.e. all the reads sequenced from a single gene are identical (therefore *X_ijr:k_* = *X_ij·:k_*, for all *r*). The sum of squares for genes within pools reads:

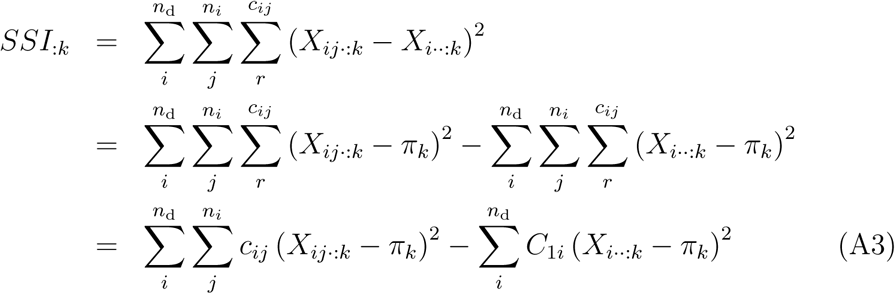

where *π_k_* is the expectation of the frequency of allele *k* over independent replicates of the evolutionary process, and 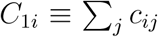 is the total number of observed reads in the *i*th pool. Likewise, the sum of squares for genes between pools reads:

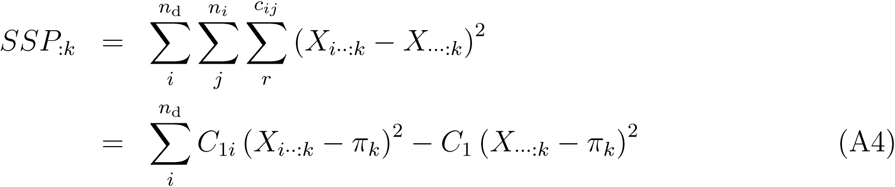

where 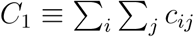 is the total number of observed reads in the full sample. These sums can be expressed as functions of the average frequency of reads of type *k* for individual 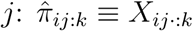, of the average frequency of reads of type *k* within the *i*th pool: 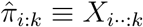, and of the average frequency of reads of type *k* in the full sample: 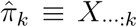. Note that from the definition of 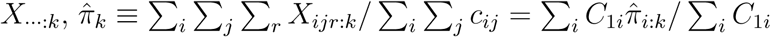 is the weighted average of the sample frequencies with weights equal to the pool coverage. Our approach is therefore equivalent to the weighted analysis-of-variance in Cockerham (1973) (see also Weir and Cockerham 1984; Weir 1996; Weir and Hill 2002; Rousset 2007; Weir and Goudet 2017). Then, developing the square in the first term in the right-hand side of Equation A3, we get:

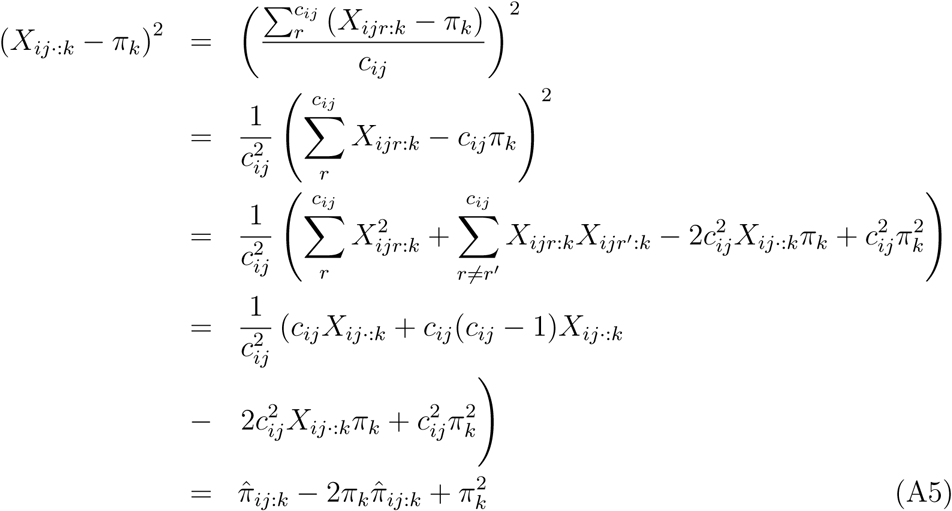

The sums of squares also depend on the unobserved frequency of pairs of genes sampled in the *i*th pool that are both of type *k*, i.e. the probability of identity in state (IIS) for allele *k*, for two distinct genes in the *i*th pool: 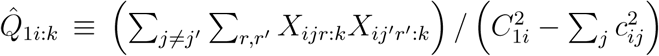. Then, developing the square in the second term in the right-hand side of Equation A3, we get:

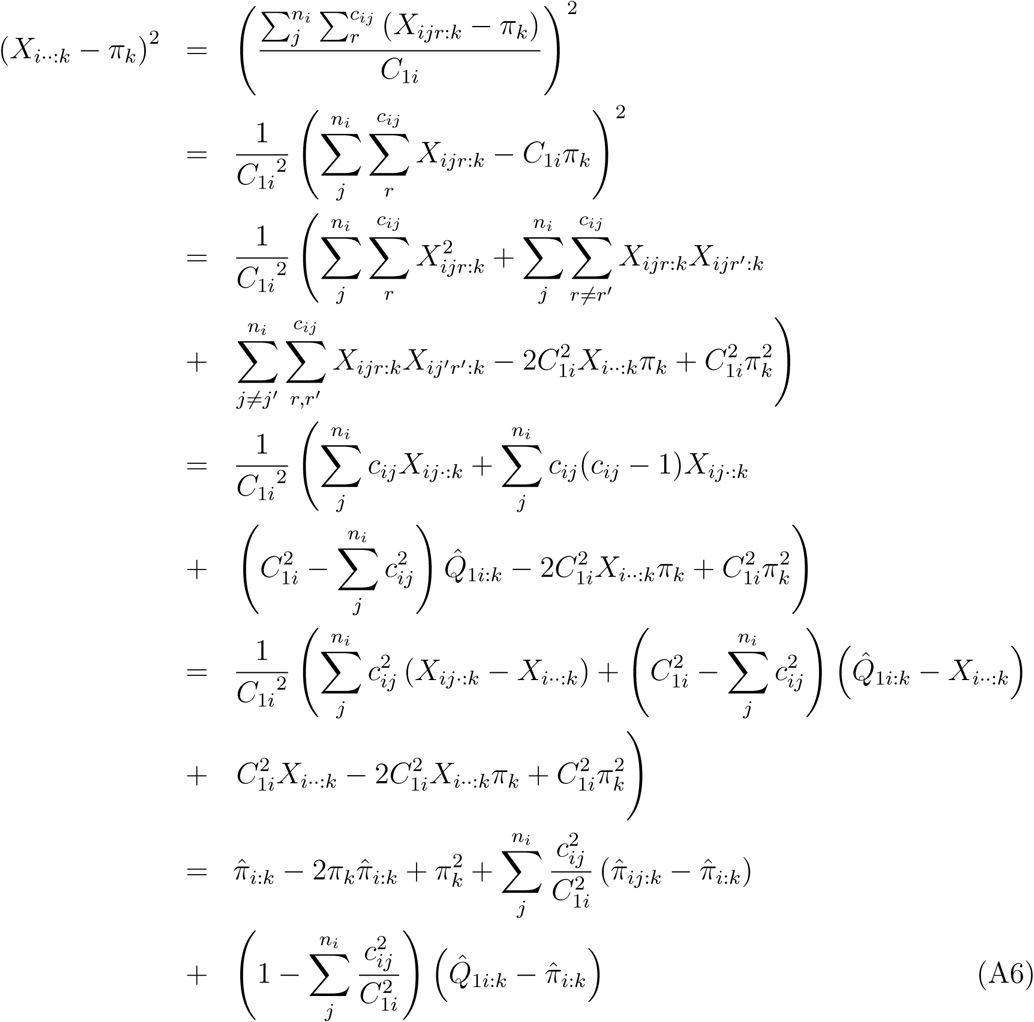

Last, the sums of squares depend on the unobserved frequency of pairs of genes sampled in the same pool that are both of type *k*, i.e. the IIS probability for allele *k* for two distinct genes in the same pool: 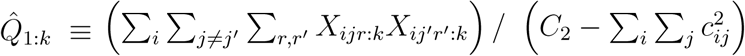, and of the unobserved frequency of pairs of genes sampled in different pools that are both of type 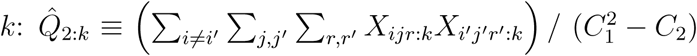, where 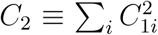.

Developing the second term in the right-hand side of Equation A4, we get:

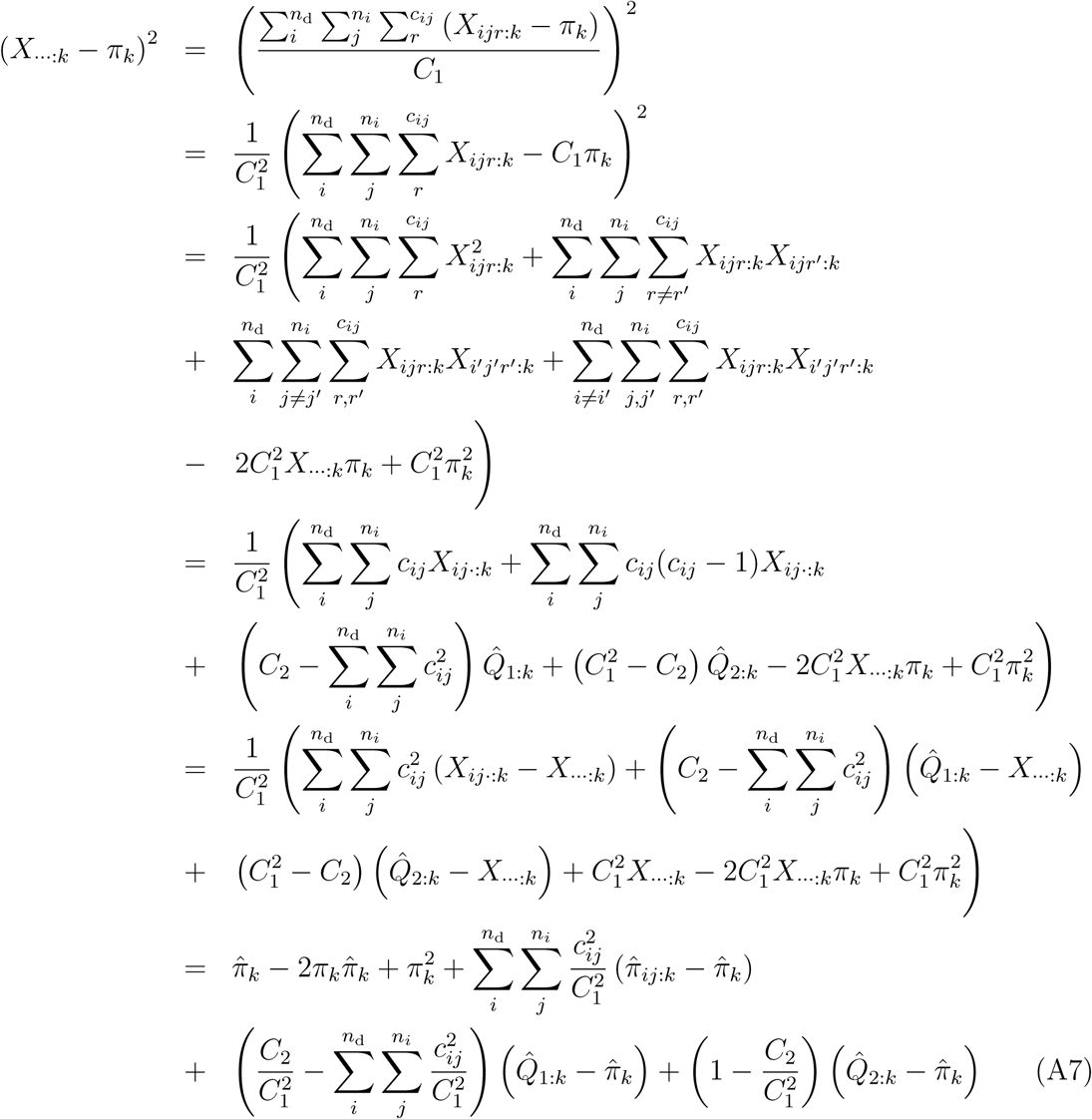

Hence, developing the first term in the right-hand side of Equation A3 using Equation A5, we have:

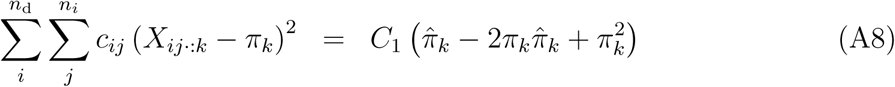

Likewise, developing the second term in the right-hand side of Equation A3 using Equation A6, we get:

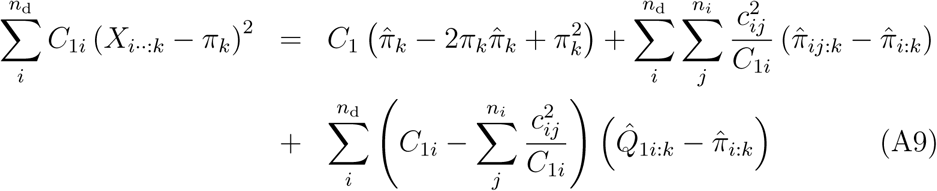

Last, developing the second term in the right-hand side of Equation A4 using Equation A7, we get:

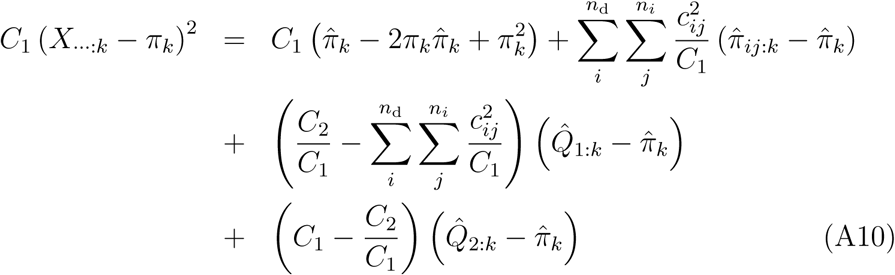

Then, from Equations A3, A8 and A9:

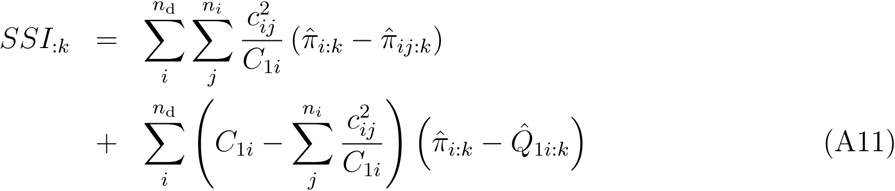

and from Equations A4, A9 and A10:

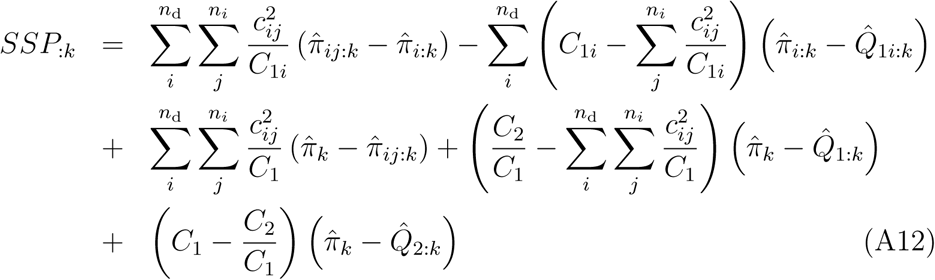

Taking expectation over all possible samples from 864 all replicate populations sharing the same evolutionary history, we get from Equation A11:

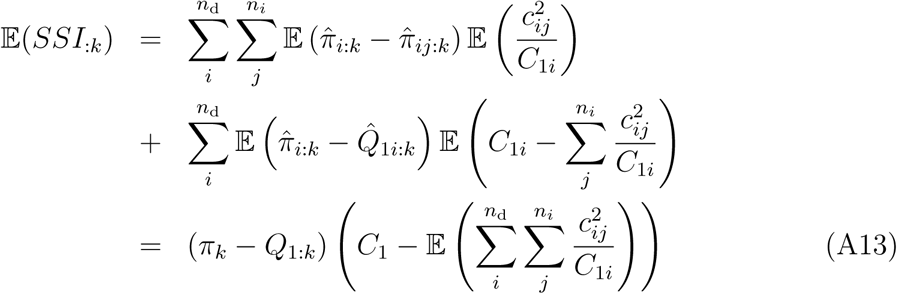

where *Q*_1:_*_k_* is the expected IIS probability that two genes in the same pool are both of type *k*. Likewise, from Equation A12:

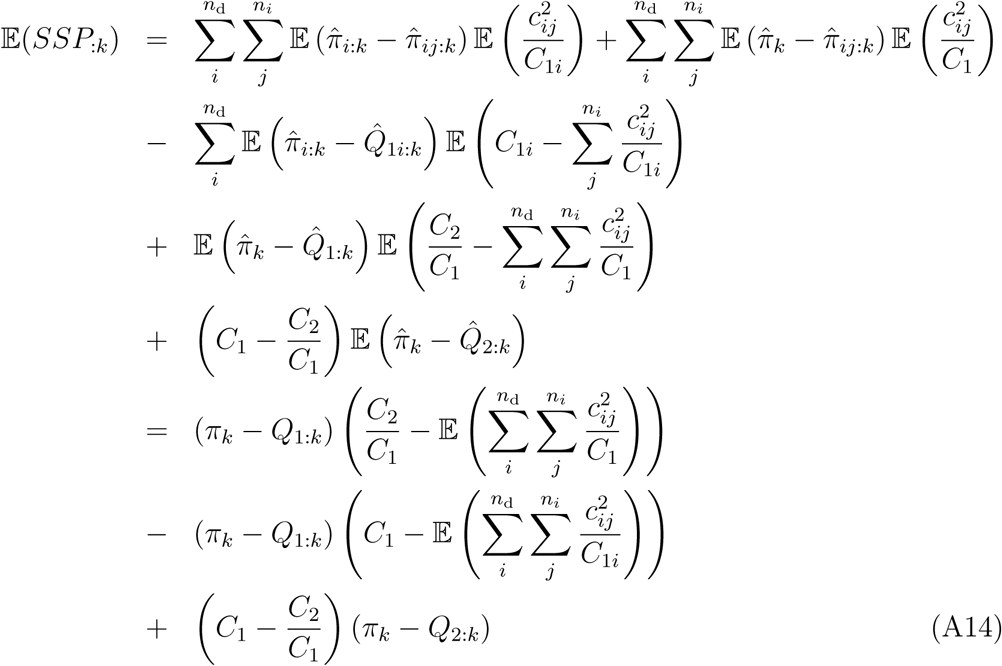

where *Q*_2:_*_k_* is the expected IIS probability that two genes from different pools are both of type *k*. Note that the expected sums 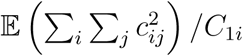 and 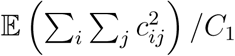 in Equations A13 and A14 depend on the latent variable *c_ij_*, that cannot be directly observed from the data. Therefore, we must make an assumption on the distribution of the *c_ij_*’s to proceed. In the following, we assume that for each pool *i*, *c_ij_* follows a multinomial distribution with parameter *C*_1_*_i_* (the number of trials, i.e. the total number of reads in the *i*th pool) and probabilities (1/*n_i_*,…, 1/*n_i_*) for the *n_i_* individuals in the pool. Then:

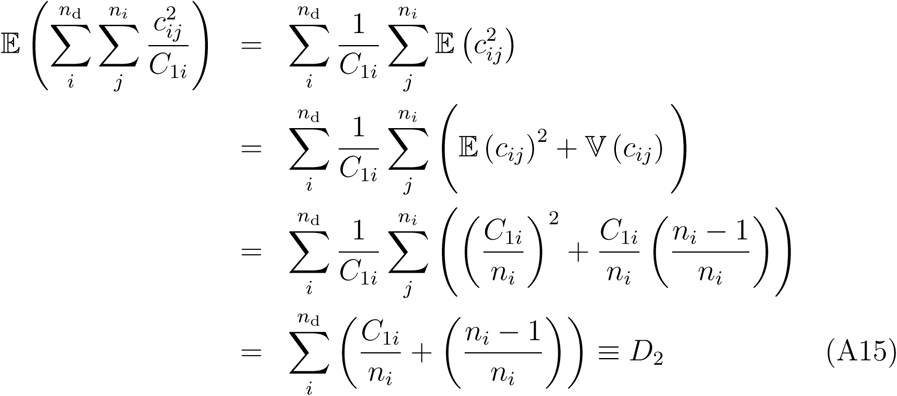

and:

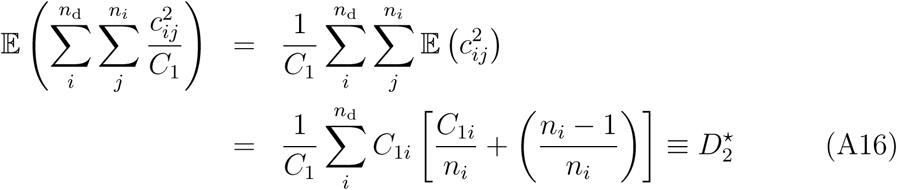

Hence, from Equations A13 and A15, we have:

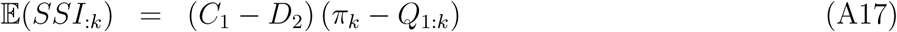

and from Equations A14 and A16:

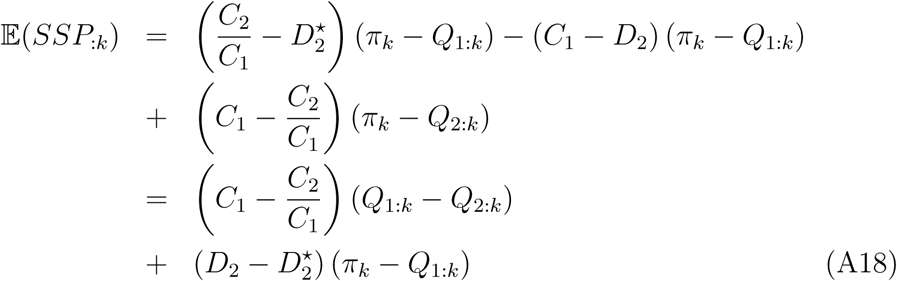

Summing over alleles, we get the following expressions for the expected sums of squares for genes between individuals within pools:

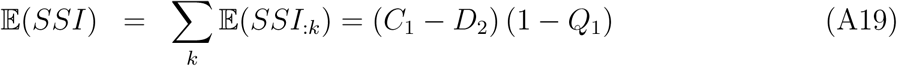

and for genes between individuals from different pools:

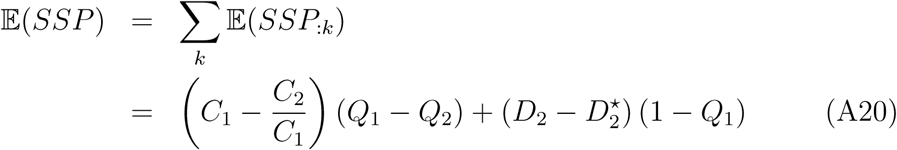

Rearranging Equations A19-A20, we get:

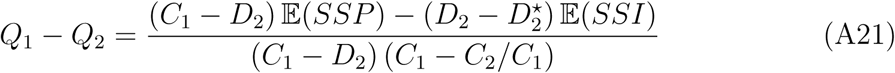

and:

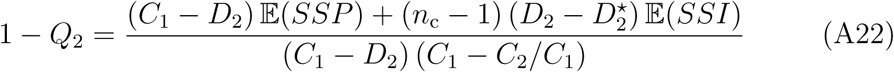

where 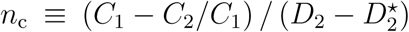. Let *MSI* ≡ *SSI*/ (*C*_1_ − *D*_2_) and 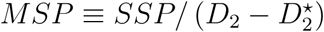. Then, rearranging Equations A21-A22, we get:

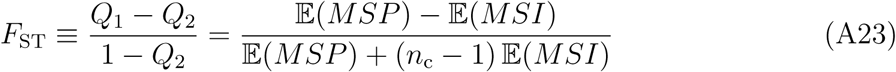

which yields the method-of-moments estimator:

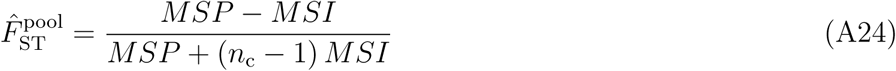

Since *SSI* (Equation A3) and *SSP* (Equation A4) may be rewritten in terms of sample frequencies as:

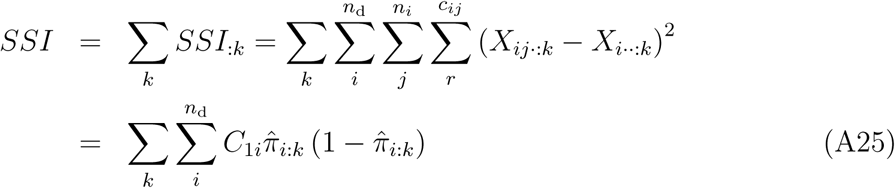

and:

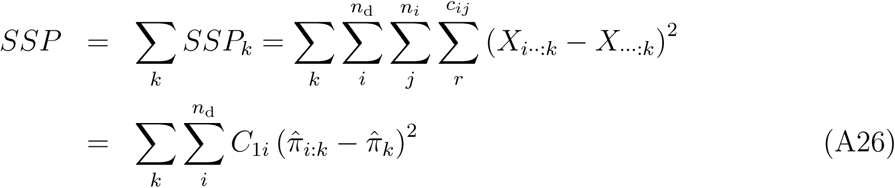

our estimator then takes the form:

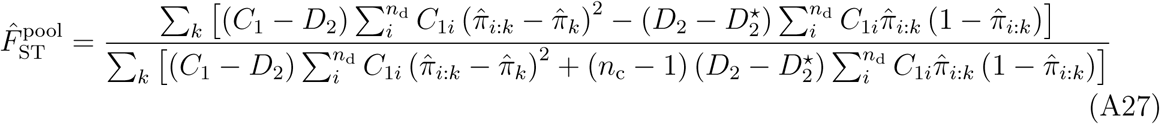

The estimator in Equation A24 can also be expressed as a function of the frequencies of identical pairs of genes 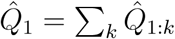 and 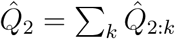, as:

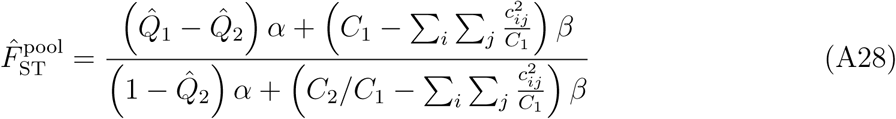

where:

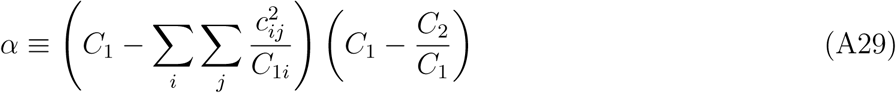

and:

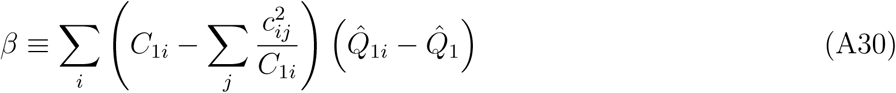

If we take the limit case where the number of sequenced reads per gene is constant, i.e. if *C*_1_*i* = *C*, for all *i* ∈ (1,…, *n*_d_), then it can be shown that Equation A28 reduces exactly to Equations 28A29-28A30 in Rousset (2007), p. 977. Furthermore, if the pools have all the same size, i.e. if *n_i_* = *n* for all *i* ∈ (1,…, *n*_d_), then 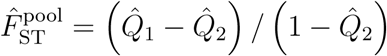.

If the pools have all the same size and if the number of reads per pool is constant, then one can also show that Equations A25-A26 reduce to:

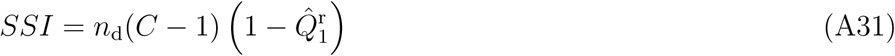

and:

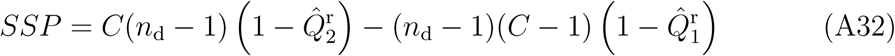

where 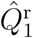 and 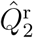 are the frequencies of identical pairs of reads within and between pools, respectively, computed by simple counting of IIS pairs. These are (unweighted) averages of the population-specific estimates 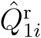 (Equation A34) and the pairwise estimates 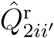 (Equation A40), respectively. Then, from Equation A24, we get:

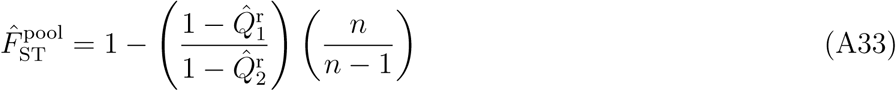

### IIS probabilities for Pool-seq data

In this Appendix, we provide unbiased estimates of IIS probabilies between pairs of genes, computed from read count data. Let 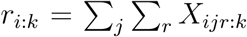 be the number of reads of type *k* in the *i*th pool. A straightforward estimate of the IIS probability between pairs of reads in the *i*th pool is given by:

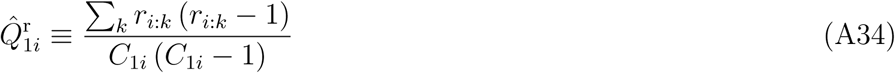

where 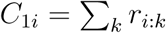. As above (see Equations A15 and A16), we assume that in each pool, the conditional distribution of the read counts *r_i:k_*, given the (unobserved) allele counts *y_i:k_*, is binomial, i.e.: *r_i:k_* | *y_i:k_* ~ Bin (*y_i:k_*/*n_i_*, *C*_1_*_i_*). The conditional expectation of the number of reads is therefore given by: 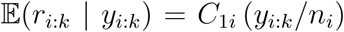, and the conditional expectation of the squared number of reads by: 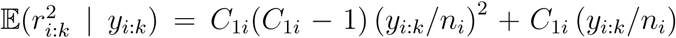. Therefore, the conditional expectation of the IIS probability between pairs of reads in the *i*th pool reads:

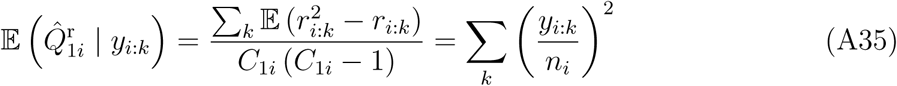

Since

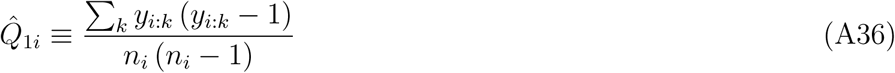

is an unbiased estimate of the IIS probability between pairs of distinct genes in the *i*th pool, Equation A35 implies that 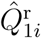 (Equation A34) is a biased estimate of that quantity (i.e., the IIS probability between pairs of reads within a pool is a biased estimate of the IIS probability between pairs of distinct genes in that pool). This is so, because the former confounds pairs of reads that are identical because they were sequenced from a single gene copy, from pairs of reads (from distinct gene copies) that are identical because they share a common ancestor. However, inspection of Equation A35 suggests that an unbiased estimate of 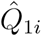 may be given by:

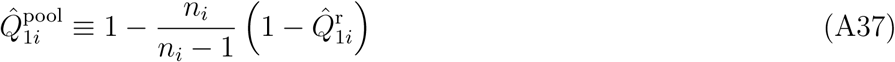

Taking expectation of Equation A37, we get indeed:

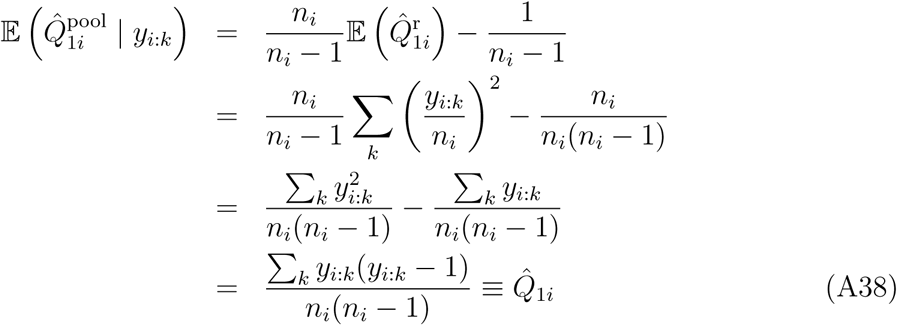

Following Weir and Goudet (2017), we define the overall IIS probability between pairs of genes within pools as the unweighted average of population-specific estimates, leading to:

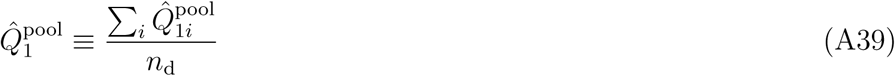

A straightforward estimate of the IIS probability between pairs of reads taken in different pools *i* and *i′* is given by:

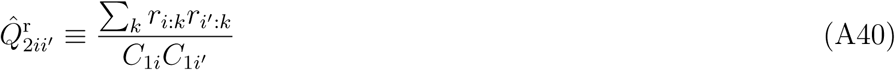

Since we assume that pools are conditionally independent, taking expectation gives:

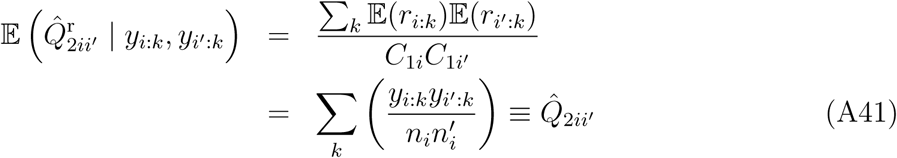

Therefore, the IIS probability between pairs of reads sampled in different pools is an unbiased estimate of the IIS probability between pairs of genes in these pools, and an unbiased estimate of the IIS probabilitiy of genes sampled from different pools is given by:

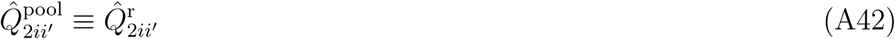

As above, we define the overall IIS probability between pairs of genes sampled from different pools as the unweighted average of pairwise estimates, i.e.:

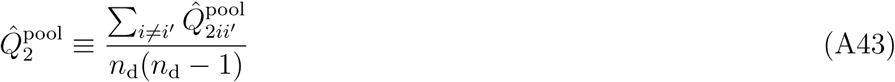

We can then derive an IIS-based estimator of *F*_ST_, as:

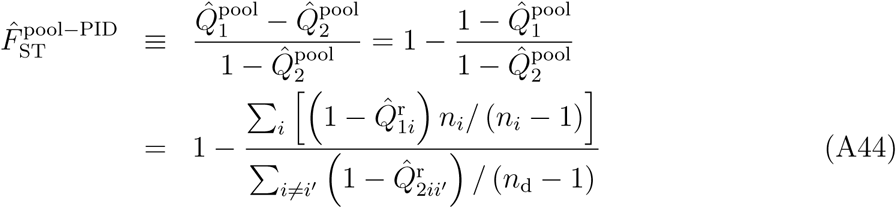

which, to the extent that we may take the expectation of a ratio to be the ratio of expectations, is unbiased. If the pools have all the same size (i.e., if *n_i_* = *n* for all *i*), then Equation A44 reduces to:

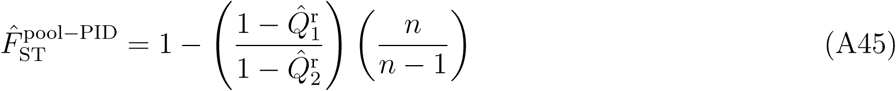

where 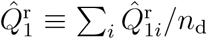 and 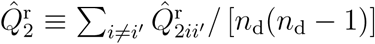. Note that Equation A45 is strictly identical to Equation A33. Therefore, if the pools have all the same size and if the number of reads per pool is constant, the analysis-of-variance estimator 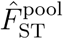 is strictly equivalent to the estimator 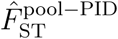 based on the computation of IIS probabilities between pairs of reads, with appropriate bias correction (see Equation A37). This echoes the derivations by Rousset (2007) for Ind-seq data, who showed that the analysis-of-variance approach (Weir and Cockerham 1984) and the simple strategy of estimating IIS probabilities by counting identical pairs of genes provides identical estimates when sample sizes are equal (see also Cockerham and Weir 1987; Karlsson et al. 2007).

Alternatively, the overall IIS probability between pairs of genes within pools may be defined as the weighted average of population-specific estimates, with weights equal to the number of pairs of genes in each pool (see Rousset 2007), i.e.:

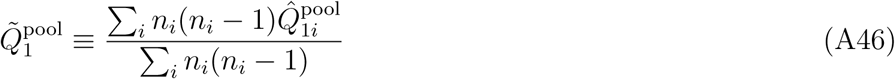

Likewise, the overall IIS probability between pairs of genes sampled from different pools may be defined as the weighted average of pairwise estimates, with weights equal to the number of pairs of genes sampled between pools, i.e:

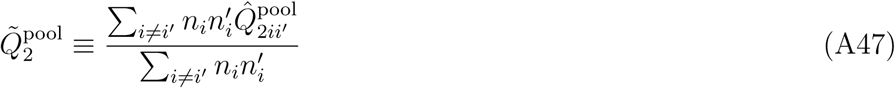

We can then derive an IIS-based estimator of *F*_ST_, using weighted IIS probabilities, as:

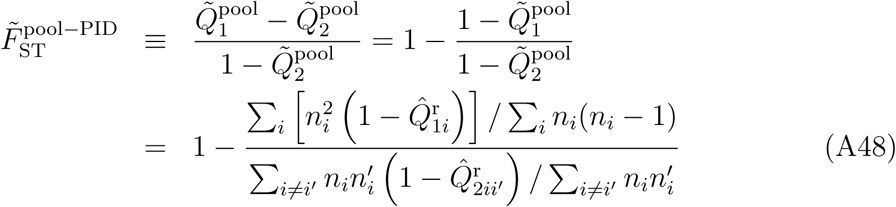

If the pools have all the same size (i.e., if *n_i_* = *n* for all *i*), then Equation A48 reduces to Equation A45, and 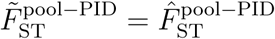. With unbalanced samples, simulation analyses show that 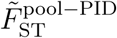 has larger bias and variance than 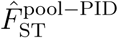, in particular for low levels of differentiation (see Figure S4).

**Table S1.** Comparison of pairwise *F*_ST_ estimates. *F*_ST_ was estimated for various conditions of expected *F*_ST_, pool size (*n*) and coverage (Cov.). For Pool-seq data, we computed our estimator 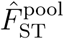 (Equation 13). The mean (RMSE) over 50 independent replicates of the `ms` simulations are provided for a single pair of populations. For comparison, we computed WC_84_ from allele count data inferred from individual genotypes (Ind-seq).

| $F_{ST}$ | $n$ | Pool-seq | | Ind-seq |
| --- | --- | --- | --- | --- |
| | | Cov. | $\hat{F}_{ST}^{pool}$ | WC<sub>84</sub> |
| 0.05 | 10 | 20× | 0.051 (0.004) |  |
| 0.05 | 10 | 50× | 0.051 (0.004) | 0.051 (0.003) |
| 0.05 | 10 | 100× | 0.051 (0.003) |  |
| 0.05 | 100 | 20× | 0.051 (0.003) |  |
| 0.05 | 100 | 50× | 0.051 (0.003) | 0.051 (0.002) |
| 0.05 | 100 | 100× | 0.051 (0.002) |  |
| 0.20 | 10 | 20× | 0.203 (0.007) |  |
| 0.20 | 10 | 50× | 0.202 (0.006) | 0.202 (0.007) |
| 0.20 | 10 | 100× | 0.201 (0.006) |  |
| 0.20 | 100 | 20× | 0.201 (0.006) |  |
| 0.20 | 100 | 50× | 0.201 (0.006) | 0.201 (0.005) |
| 0.20 | 100 | 100× | 0.202 (0.005) |  |

**Table S2.** Effect of unequal sampling on pairwise *F*_ST_ estimates. Pairwise *F*_ST_ was estimated for various conditions of expected *F*_ST_ and coverage (Cov.). The pool size (*n*) was variable across demes, with haploid sample size *n* drawn independently for each deme from a Gaussian distribution with mean 100 and standard deviation 30; *n* was rounded up to the nearest integer, with min. 20 and max. 300 haploids per deme. For Pool-seq data, we computed our estimator 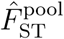 (Equation 13). The mean (RMSE) over 50 independent replicates of the ms simulations are provided, for a single pair of populations. For comparison, we computed WC_84_ (Weir and Cockerham 1984) from allele count data inferred from individual genotypes (Ind-seq).

| $F_{ST}$ | $n$ | Pool-seq | | Ind-seq |
| --- | --- | --- | --- | --- |
| | | Cov. | $\hat{F}_{ST}^{pool}$ | WC<sub>84</sub> |
| 0.05 | $\mathcal{N}(100, 30)$ | 20× | 0.051 (0.003) | |
| 0.05 | $\mathcal{N}(100, 30)$ | 50× | 0.052 (0.003) | 0.051 (0.002) |
| 0.05 | $\mathcal{N}(100, 30)$ | 100× | 0.051 (0.002) | |
| 0.20 | $\mathcal{N}(100, 30)$ | 20× | 0.202 (0.007) | |
| 0.20 | $\mathcal{N}(100, 30)$ | 50× | 0.202 (0.006) | 0.202 (0.006) |
| 0.20 | $\mathcal{N}(100, 30)$ | 100× | 0.202 (0.006) | |

**Table S3.**
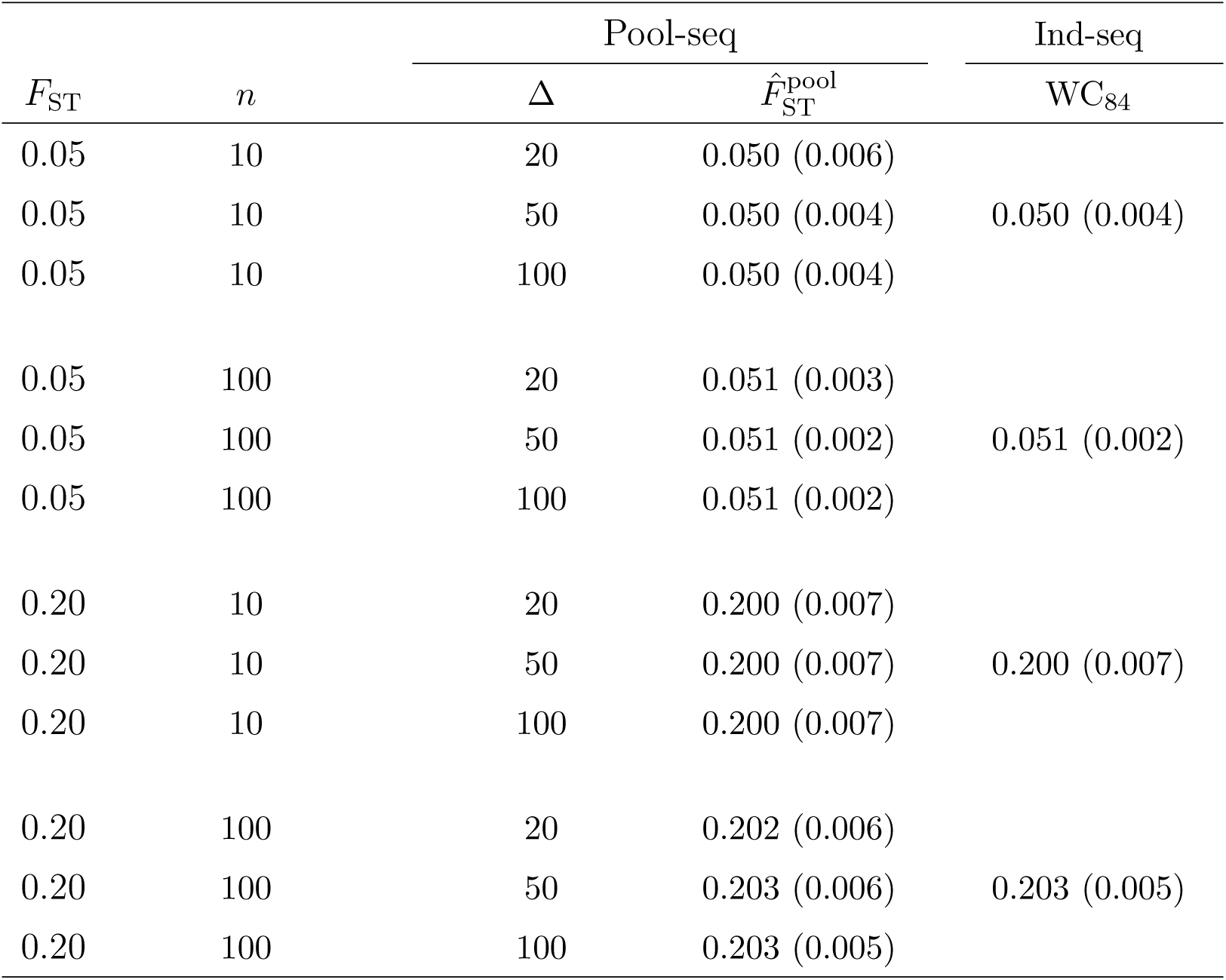
Effect of variable coverage on pairwise F_ST_ estimates. Pairwise *F*_ST_ was estimated for various conditions of expected *F*_ST_ and pool size (*n*). The coverage (*δ_i_*) was varying across demes and loci, with *δ_i_* ~ Pois(∆). For Pool-seq data, we computed our estimator 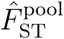 (Equation 13). The mean (RMSE) over 50 independent replicates of the `ms` simulations are provided, for a single pair of populations. For comparison, we computed WC_84_ from allele count data inferred from individual genotypes (Ind-seq).

| $F_{ST}$ | $n$ | Pool-seq | | Ind-seq |
| --- | --- | --- | --- | --- |
| | | $\Delta$ | $\hat{F}_{ST}^{pool}$ | WC<sub>84</sub> |
| 0.05 | 10 | 20 | 0.050 (0.006) |  |
| 0.05 | 10 | 50 | 0.050 (0.004) | 0.050 (0.004) |
| 0.05 | 10 | 100 | 0.050 (0.004) |  |
| 0.05 | 100 | 20 | 0.051 (0.003) |  |
| 0.05 | 100 | 50 | 0.051 (0.002) | 0.051 (0.002) |
| 0.05 | 100 | 100 | 0.051 (0.002) |  |
| 0.20 | 10 | 20 | 0.200 (0.007) |  |
| 0.20 | 10 | 50 | 0.200 (0.007) | 0.200 (0.007) |
| 0.20 | 10 | 100 | 0.200 (0.007) |  |
| 0.20 | 100 | 20 | 0.202 (0.006) |  |
| 0.20 | 100 | 50 | 0.203 (0.006) | 0.203 (0.005) |
| 0.20 | 100 | 100 | 0.203 (0.005) |  |

**Figure S1.**
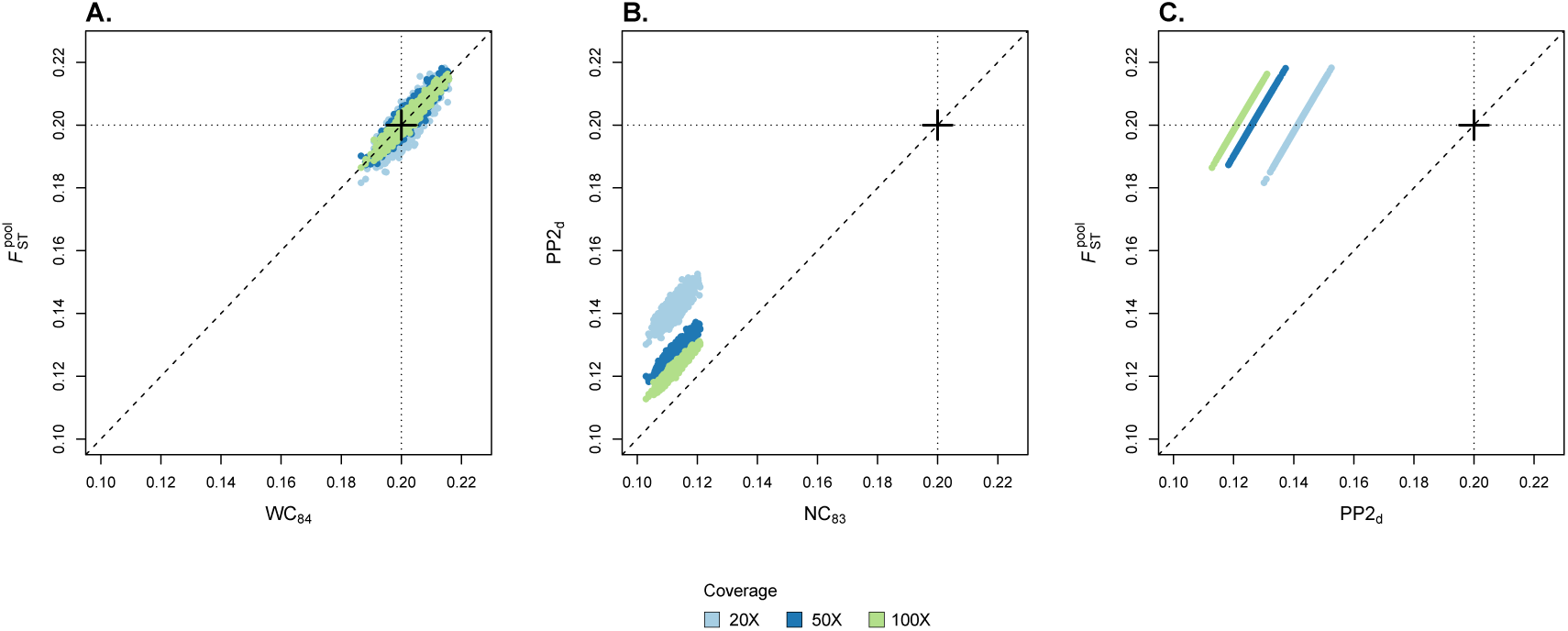
Pairwise estimators of *F*_ST_. A. Multilocus estimates 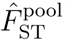 (computed using Equation 13) as a function of WC_84_ estimates computed from allele count data inferred from individual genotypes. B. Multilocus estimates PP2_d_, as a function of NC_83_ estimates computed from allele count data inferred from individual genotypes. C. Multilocus estimates 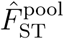 as a function of PP2_d_ estimates. In each graph, the dots represent multilocus estimates of *F*_ST_ across all pairs of subpopulations from an 8-island model, and across 50 replicate `ms` simulations. We specified the migration rate corresponding to *F*_ST_ = 0.20. The size of each pool was fixed to 100. The results are shown for different coverages (20X, 50X and 100X). The cross indicates the simulated value of the parameter *F*_ST_.

**Figure S2.**
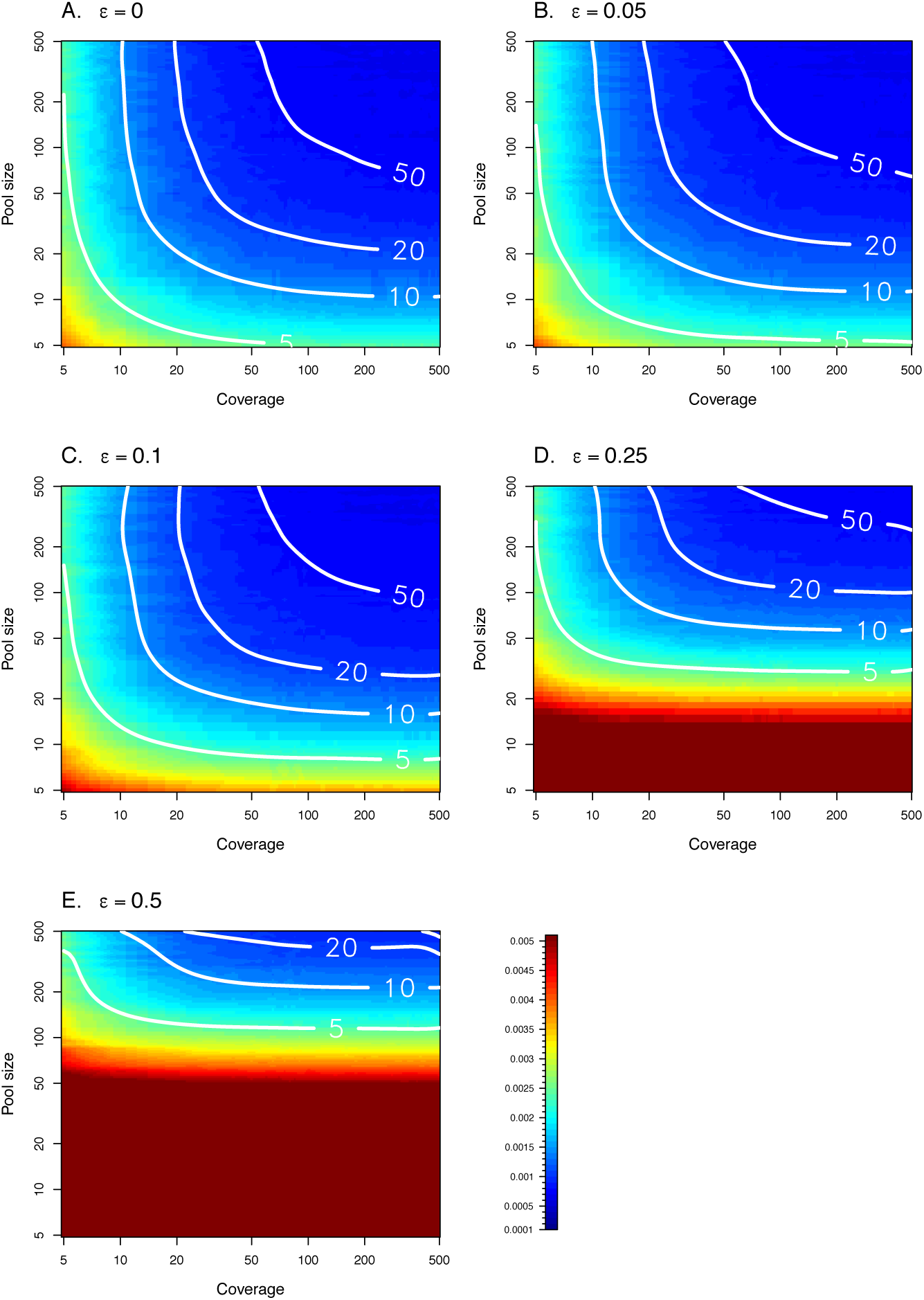
Root mean squared error (RMSE) of *F*_ST_ estimates for a wide range of pool sizes and coverage, with experimental error rate *∊* varying from 0 to 0.5 (A-E). Each density plot gives the RMSE of our estimator 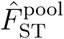, using simple linear interpolation from a set of 44 × 44 pairs of pool size and coverage values. For each pool size and coverage, 500 replicates of 5,000 markers were simulated. Plain white isolines represent the RMSE of the WC_84_ estimator computed from Ind-seq data, for various sample sizes (*n* = 5, 10, 20, and 50). Each isoline was fitted using a thin plate spline regression with smoothing parameter λ = 0.005, implemented in the `fields` package for R (Nychka et al. 2017).

**Figure S3.**
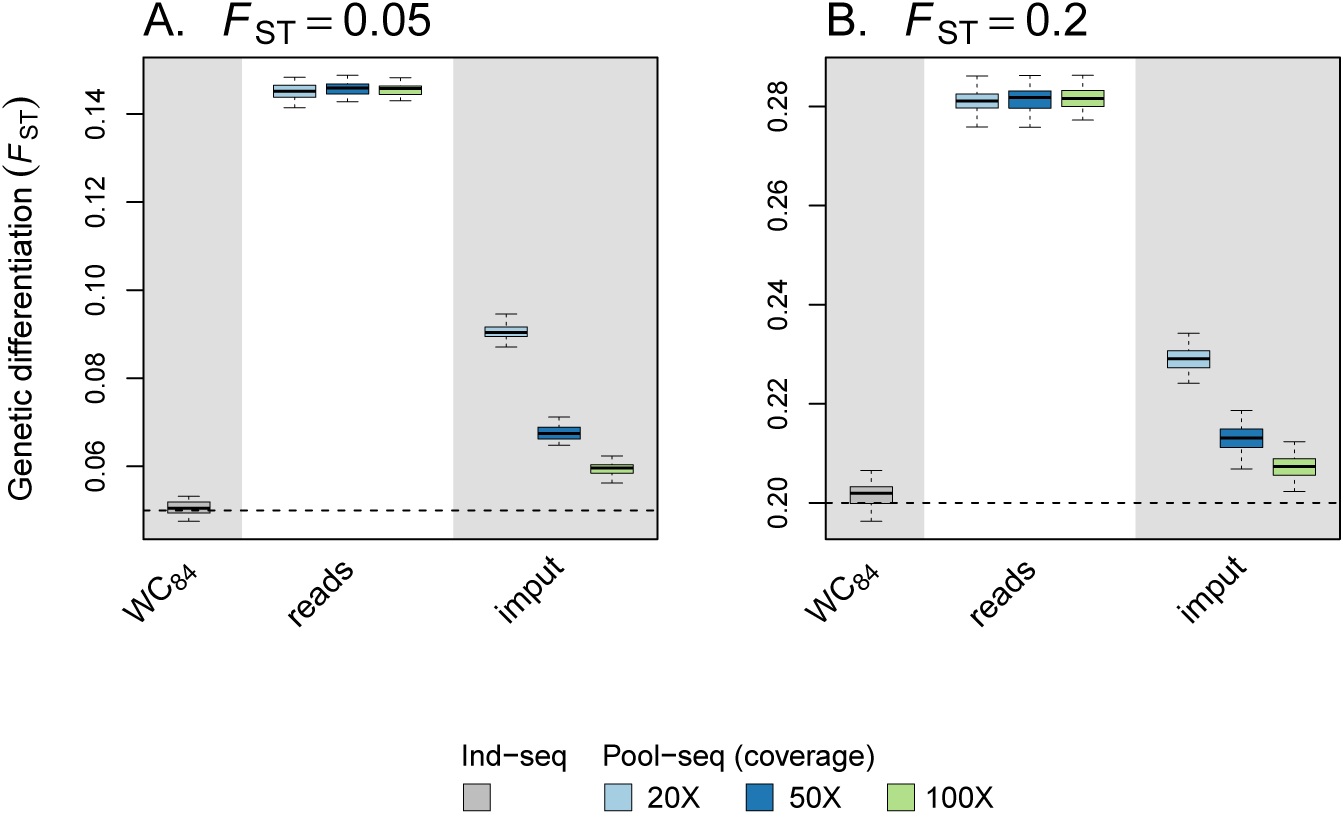
Global estimators of *F*_ST_. We considered one estimator based on allele count data inferred from individual genotypes (Ind-seq): WC_84_. For pooled data, we computed *F*_ST_ using the WC_84_ estimator: (i) directly from read counts, as if they were allele counts (“reads”); (ii) from allele counts imputed by maximum-likelihood (“imput”), as in Leblois et al. (2018). Each boxplot represents the distribution of multilocus *F*_ST_ estimates across all demes comparisons in an 8-island model, and across 50 independent replicates of the `ms` simulations. We used two migration rates, corresponding to *F*_ST_ = 0.05 (A) or *F*_ST_ = 0.20 (B). The size of each pool was fixed to 10. For Pool-seq data, we show the results for different coverages (20X, 50X and 100X). In each graph, the dashed line indicates the simulated value of *F*_ST_.

**Figure S4.**
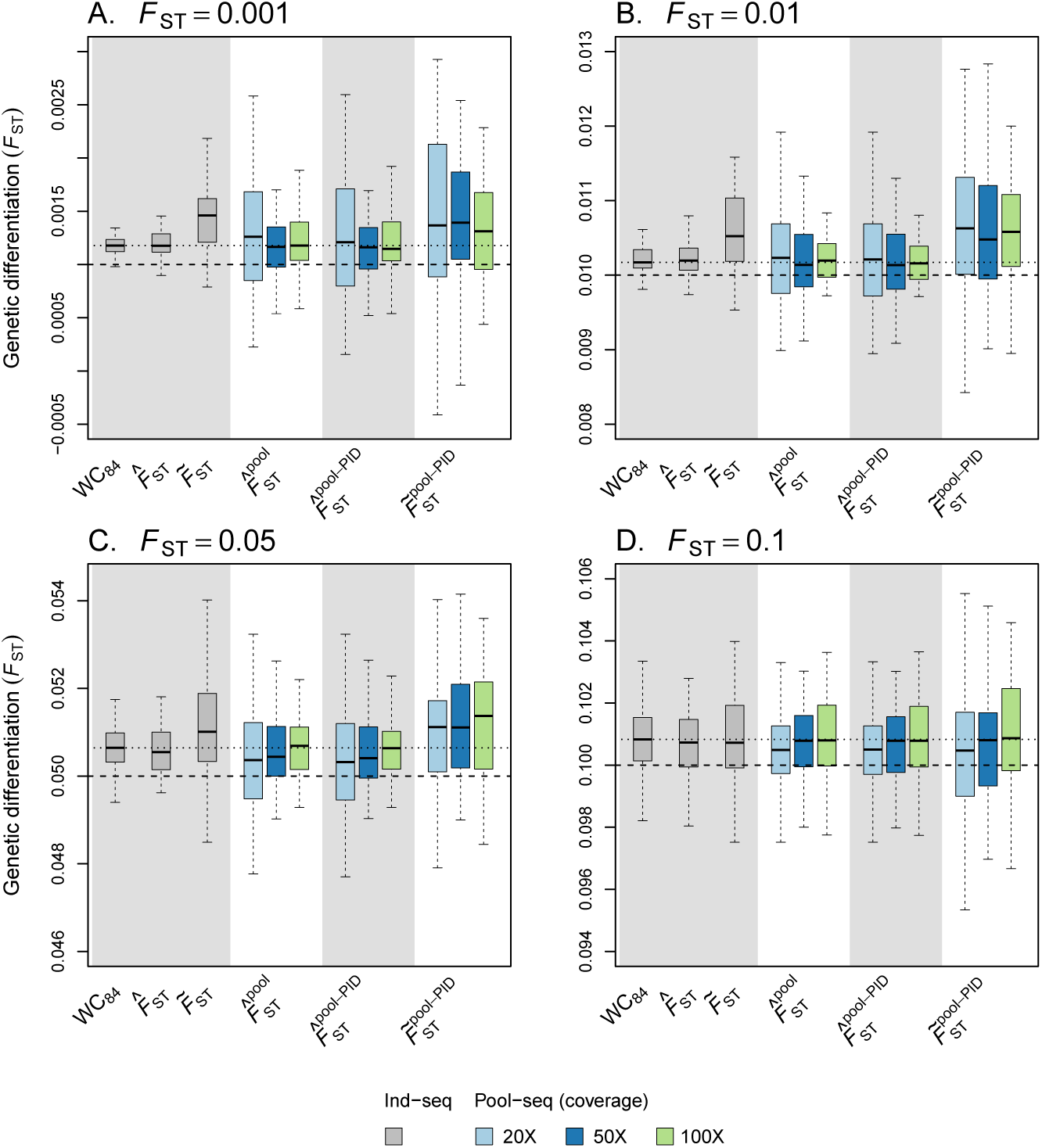
Precision and accuracy of alternative estimators of *F*_ST_ with varying pool size, for various levels of differentiation (A-D). The haploid pool size *n* drawn independently for each deme from a Gaussian distribution with mean 100 and standard deviation 30; *n* was rounded up to the nearest integer, with min. 20 and max. 300 haploids per deme. We considered three estimators based on allele count data inferred from individual genotypes (Ind-seq): WC_84_, 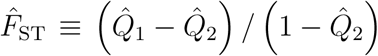 (where 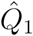 and 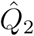 are the weighted frequencies of identical pairs of genes within and between subpopulations, respectively, with weights equal to the number of pairs of genes) and 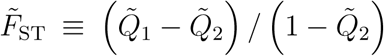 (where 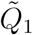 and 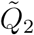 are the unweighted frequencies of identical pairs of genes within and between subpopulations, respectively. For Pool-seq data, we considered the estimators 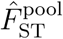 (Equation 12), 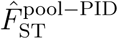 (Equation A44) and 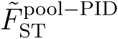 (Equation A45). Each boxplot represents the distribution of multilocus *F*_ST_ across 50 independent replicates of the `ms` simulations. For Pool-seq data, we show the results for different coverages (20X, 50X and 100X). In each graph, the dashed line indicates the simulated value of *F*_ST_ and the dotted line indicates the median of the distribution of WC_84_ estimates.

